# Influence of Temperate Forest Autumn Leaf Phenology on Segmentation of Tree Species from UAV Imagery Using Deep Learning

**DOI:** 10.1101/2023.08.03.548604

**Authors:** Myriam Cloutier, Mickaël Germain, Etienne Laliberté

## Abstract

Remote sensing of forests has become increasingly accessible with the use of unoccupied aerial vehicles (UAV), along with deep learning, allowing for repeated high-resolution imagery and the capturing of phenological changes at larger spatial and temporal scales. In temperate forests during autumn, leaf senescence occurs when leaves change colour and drop. However, the influence of leaf senescence in temperate forests on tree species segmentation using a Convolutional Neural Network (CNN) has not yet been evaluated. Here, we acquired high-resolution UAV imagery over a temperate forest in Quebec, Canada on seven occasions between May and October 2021. We segmented and labelled 23,000 tree crowns from 14 different classes to train and validate a CNN for each imagery acquisition. The CNN-based segmentation showed the highest F1-score (0.72) at the start of leaf colouring in early September and the lowest F1-score (0.61) at peak fall colouring in early October. The timing of the events occurring during senescence, such as leaf colouring and leaf fall, varied substantially between and within species and according to environmental conditions, leading to higher variability in the remotely sensed signal. Deciduous and evergreen tree species that presented distinctive and less temporally-variable traits between individuals were better classified. While tree segmentation in a heterogenous forest remains challenging, UAV imagery and deep learning show high potential in mapping tree species. Our results from a temperate forest with strong leaf colour changes during autumn senescence show that the best performance for tree species segmentation occurs at the onset of this colour change.

## 1. Introduction

Monitoring vegetation and detecting changes in biodiversity is critical for understanding the effects of climate change on plant species distribution and ecosystem health. Improved knowledge and data on forest dynamics are needed for efficient and sustainable management of these ecosystems (Fassnacht et al., 2016; Goodbody et al., 2019; Lausch et al., 2017). To acquire these data, spatially explicit data on forest stands need to be produced periodically and ideally at the individual tree level (Korpela & Tokola, 2006; Natesan et al., 2020). However, identifying individual tree species can be difficult depending on the complexity of the forest and the variability of the traits of the target species (Fassnacht et al., 2016; Natesan et al., 2020; Zhang et al., 2022). Moreover, traditional methods used for field surveys in forests are limited to small spatial extents, are time-consuming and labour-intensive, and can be very expensive, especially in remote regions. These factors make it challenging to conduct repeated forest surveys (Beloiu et al., 2023; Goodbody et al., 2019; Veras et al., 2022).

Remote sensing, specifically from unoccupied aerial vehicles (UAVs), has been identified in recent years as a tool capable of addressing these challenges and benefiting research in plant ecology and forestry (Cavender-Bares et al., 2022; de Lima et al., 2022; Lechner et al., 2020). Forests that require monitoring and management will increasingly benefit from remote sensing with UAVs (Dainelli et al., 2021). The many advantages of UAVs compared to on-the-ground field surveys have led to an increase in their use in ecology, resulting in an increase in available data, as well as higher diversity of these data (e.g. spatial, temporal and spectral; Kattenborn et al., 2021). UAVs are flexible, increasingly accessible, and inexpensive, enabling the acquisition of multi-temporal data for tree species in heterogeneous forests (Klosterman & Richardson, 2017; Komárek, 2020; Veras et al., 2022). UAVs also have the potential to acquire very high spatial resolution RGB imagery, making it possible to see detailed characteristics in the tree crowns, such as leaf and canopy shape and branching patterns (Fassnacht et al., 2016; Kattenborn et al., 2019; Natesan et al., 2020). High spatial resolution is necessary for these patterns to be visible in the imagery and can thus help to identify tree species using recent computer vision approaches (Schiefer et al., 2020).

High-resolution UAV data require methods that can extract the complex features present in the imagery. Recently developed deep learning algorithms have been used to process the imagery by efficiently extracting the features present in forest canopies to map vegetation with high accuracy (Brodrick et al., 2019; Katal et al., 2022; Veras et al., 2022). Convolutional Neural Networks (CNN), a type of deep learning algorithms, are capable of extracting complex spatial information about objects of interest from the imagery (Kattenborn et al., 2021), as well as the spatial context surrounding object of interest (Korznikov et al., 2021). When acquired repeatedly across a growing season, UAV imagery data combined with deep learning can also leverage species-specific phenological patterns (Veras et al., 2022). CNNs were specifically developed to analyze spatial patterns, making them highly effective when using high spatial resolution remote sensing data for spatial prediction problems by leveraging higher detail in the imagery (Fricker et al., 2019; Kattenborn et al., 2019; Schiefer et al., 2020). Other advantages of using CNNs on remote sensing RGB imagery is the high classification accuracy, which can approach and even surpass human vision in certain situations, but that can be rapidly applied to a large extent in an automatic fashion (Brodrick et al., 2019). This can allow for rapid mapping of trees at the species-level over large extents, provided that sufficient spatially explicit reference data is available (Kattenborn et al., 2019; Weinstein et al., 2019).

Despite the increasing accessibility of UAVs and deep learning methods, identifying tree species from UAV imagery remains a challenge in natural, heterogeneous forests (Fassnacht et al., 2016; Zhang et al., 2022). Some studies have successfully mapped tree species in heterogeneous forests in temperate regions (Beloiu et al., 2023; Gan et al., 2023; Schiefer et al., 2020) and tropical regions (Braga et al., 2020; Veras et al., 2022) using a combination of high-resolution RGB imagery from UAVs with deep learning. However, challenges remain in complex mixed-species forest canopies due to overlapping crowns and diverse crown structure, especially when it comes to individual tree crown segmentation (Zhang et al., 2022). To potentially overcome this challenge, many researchers have suggested the use of multi-temporal imagery (Hill et al., 2010; Veras et al., 2022; Yang et al., 2017). However, these studies include different phenological events in the training dataset without knowing which aspects of the vegetation may influence the accuracy (Kattenborn et al., 2021; Yang et al., 2019; Zhang et al., 2018), limiting our efficiency in leveraging phenological information to classify tree species.

Phenology is defined as the timing of seasonal events. Foliar phenology in particular can be used to differentiate between tree species and increase classification accuracy (Fassnacht et al., 2016; Nagendra et al., 2013), as leaves are the most prominent organs that are captured in aerial imagery. Consequently, foliar phenological characteristics of tree species can be particularly useful for tree species classification in complex tree canopies. When training a CNN, the amount of reference data needed can be reduced by harnessing these species-specific phenological patterns that increase the contrast between classes of interest (Kattenborn et al., 2021; Veras et al., 2022). Specifically, in autumn, leaf senescence is the phenological event during which leaves age due to a decline in function which is triggered by a systemic response in the plant, causing leaves to change colours before falling (Gallinat et al., 2015; Zani et al., 2020). However, when compared to research focusing on spring phenology, autumn phenology has received significantly less attention (Zani et al., 2020). This can be attributed to the higher complexity regarding the drivers of senescence and its unpredictable timing year-to-year (Gallinat et al., 2015; Katal et al., 2022). Additionally, leaf senescence exhibits a greater interspecies variation than for spring phenology, such as for leaf budding and leaf unfolding (Budianti et al., 2021). This makes it challenging to select an appropriate date for acquiring UAV imagery to capture leaf senescence across many tree species.

More broadly, further research is needed at a high resolution on large scales on tree autumn coloration, an important phenomenon found in temperate forests Northeastern North America, Europe and Asia (Renner & Zohner, 2019). This bright display of red, orange and yellow autumn leaf colours in these temperate regions could be affected by changing global temperatures and should be included in climate change models (Norby et al., 2003). In fact, many authors who have investigated the use of UAV and deep learning for mapping tree species in temperate forests mention a need for studies examining the effects of phenology on these methods (Beloiu et al., 2023; Gan et al., 2023; Korznikov et al., 2021; Nuijten et al., 2019). More broadly, there is a need for additional research aimed at identifying tree species traits that could help or hinder classification accuracy of a CNN model (Fassnacht et al., 2021; Onishi & Ise, 2021).

In this study, we assessed the influence of autumn phenology on the task of segmenting tree species from high-resolution UAV RGB imagery in a temperate mixed forest in Eastern Canada, using deep learning. To do so, we trained seven separate CNNs using imagery from seven UAV acquisitions over the same forest during the 2021 growing season, using a large reference dataset in which 23,000 individual tree crowns were segmented and annotated to species at the tree level. We evaluated the performance of the different CNN models for each species and each date. We hypothesized that the model trained with the imagery taken during peak autumn colour (i.e. early October) would perform the best out of the entire growing season. We expected the colour change to help differentiate between species and to highlight the contours of the crowns. We also hypothesized that tree classes that were closer to one another phylogenetically (i.e. tree species within the same genus) would show a higher degree of confusion in the predictions because of similar foliar traits. Overall, our aim was to evaluate the change in performance of a CNN in segmenting 14 classes (11 tree species, two genera and dead trees) according to the timing in the growing season, and to determine some of the characteristics of the tree species that would explain the differences in performance, specifically leaf senescence and other species-specific traits, such as colour, texture, and shape.

## 2. Methods

### 2.1 Study site

The study was conducted in a mixed temperate forest in Quebec, Canada. The study site is located within the research station of Université de Montréal in Saint-Hippolyte (45°59’17.34” N / 74° 0’20.75” W, Figure 1). The site has a diverse topography, with hills, cliffs, and valleys, bringing a range of different forest types and environmental conditions within the study area. The study area is mostly covered by unmanaged mixed temperate forest, which includes a variety of deciduous and evergreen tree species, as well as a small bog and many lakes. The area has been affected by disturbances in the last century, such as logging and medium to low intensity fires, before becoming a research station in 1965 (Savage, 2001). The most common tree species that form the canopy are *Betula papyrifera* Marshall, *Acer rubrum* L., *Populus grandidentata* Michaux and *Acer saccharum* Marshall (Savage, 2001).

**Figure 1.**
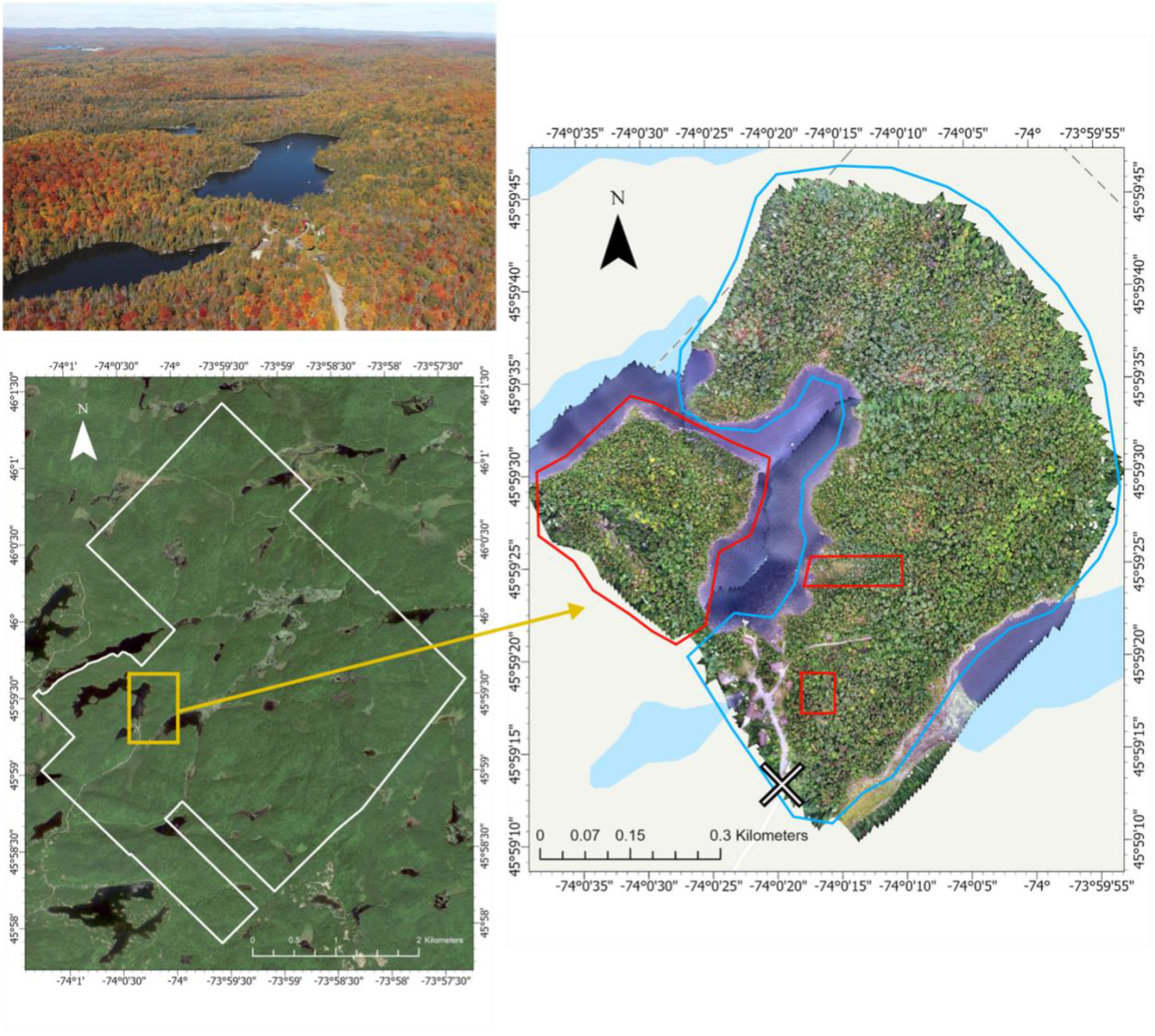
Map of the study site in Saint-Hippolyte, Quebec, Canada. The biological research station is delineated in white. The study site used for training and validation is delineated in blue and the test site is in red. The two sections used for the test dataset (smaller polygons in red within the blue polygon) were excluded from the training and validation dataset and served to assure all classes were present in both datasets. The photography at the top was taken from the position marked with an X and shows the study site. Projection: WGS84 UTM Zone 18N (EPSG:32618).

The study area was divided into a training and validation site and a test site, totalling about 44 ha. The imagery for the training and validation site was acquired using two different flights due to the drone’s maximum flight time and a third flight was done for the imagery for the test site. The three flights were taken near the middle of the day, before and after noon, to minimize changes in light conditions. Regarding the division of the area, we selected the largest area as the training site because it contained most of the tree crowns in our dataset. The test site was chosen because it was physically separated from the training site with a lake while containing enough tree crowns for inference and model performance evaluation. This separation of the training site and the test site was done to minimize spatial autocorrelation between the two datasets (Kattenborn et al., 2022).

### 2.2 Datasets

#### 2.2.1 Drone imagery

The imagery was acquired using a UAV, a DJI Phantom 4 RTK aircraft (DJI Science and Technology Co. Ltd., Shenzhen, China). This UAV is equipped with a Real Time Kinematic (RTK) GNSS receiver which is synchronized with the inertial measurement unit (IMU) and the camera, allowing for direct georeferencing of each photo (Kalacska et al., 2020; Taddia et al., 2019). The imagery was acquired 60 meters above the canopy. To maintain a constant distance between the UAV and the canopy across the study area, we used DJI’s Terrain Follow Mode using a 1-m LiDAR-derived digital surface model as our reference (Gouvernement du Québec, 2021).

The imagery was acquired seven times in 2021, at a rate of one acquisition per month from May to August 2021, and three acquisitions for September and October during the period of leaf senescence. We selected these dates to have a temporal resolution that allowed us to see the changes in foliage during the growing season, as well as the faster rate of change in leaf colour in autumn. We acquired the imagery on days during which the acquisition would not be affected by strong wind or rain. However, the light conditions varied from one imagery to the other, ranging from overcast conditions, to passing clouds, to full sun (Figure S1). The imagery with poorer light conditions (i.e., lower cloud coverage) were of good quality, so we did not adjust the images to homogenize the light conditions. To ensure the imagery was well georeferenced, we positioned 25 ground control points (GCPs) throughout the area. The coordinates of the ground control points were then measured using two Emlid Reach RS2 RTK GNSS receivers, one as a base on a geodetic marker and one as a rover.

The aerial imagery was processed using Agisoft Metashape Professional v.1.7.5 (Agisoft, St. Petersburg, Russia) to generate orthomosaics using Structure from motion photogrammetry. Three orthomosaics were made for each of the dates, according to the drone flights (two for training site and one for testing site), totalling 21 orthomosaics. These orthomosaics had a ground sampling distance between 1.81 and 2.02 cm. The protocol used to generate the orthomosaics was adapted from Over et al. (2021). We started by importing and aligning the drone photos and importing the GCPs coordinates into Metashape to precisely georeference the images. We then created a dense point cloud using the tie points created during the image alignment. We set our parameters to aggressive filtering and medium density to create the dense point cloud. This approach made it smoother and helped avoid local artefacts and noise when using the dense point cloud to build the digital elevation models (DEM), which were then used to build the orthomosaics by projecting the images onto the DEM.

#### 2.2.2 Annotations

We conducted a field survey the same summer the imagery was taken, in 2021. Using an Arrow 100 submeter GNSS receiver (EOS Positioning Systems, Inc. Terrebonne, Canada) and a Windows Surface Pro tablet, we generated a label for each tree crown on the orthomosaics in the form of a point directly in the field. This was done using ArcGIS Pro v.2.9 (ESRI Inc., Redlands, CA, USA) from which we could visualize an orthomosaic and assign the labels using a point layer. The Arrow 100, connected to the tablet, allowed us to locate ourselves on the orthomosaic to minimize labelling errors. In ArcGIS Pro, the points were then used to create polygons delineating the tree crowns individually according to the class assigned during the field survey (Figure 2). In total, we delineated 23,000 tree crowns. The parts of the imagery we did not annotate were classified as background, mainly forest floor, herbaceous vegetation, rocks, water, and buildings. This also includes trees from the classes we were interested in, but for which the crowns were not clear enough to delineate due to overlapping crowns or blurriness in the orthomosaic or that have not been annotated during the field survey.

**Figure 2.**
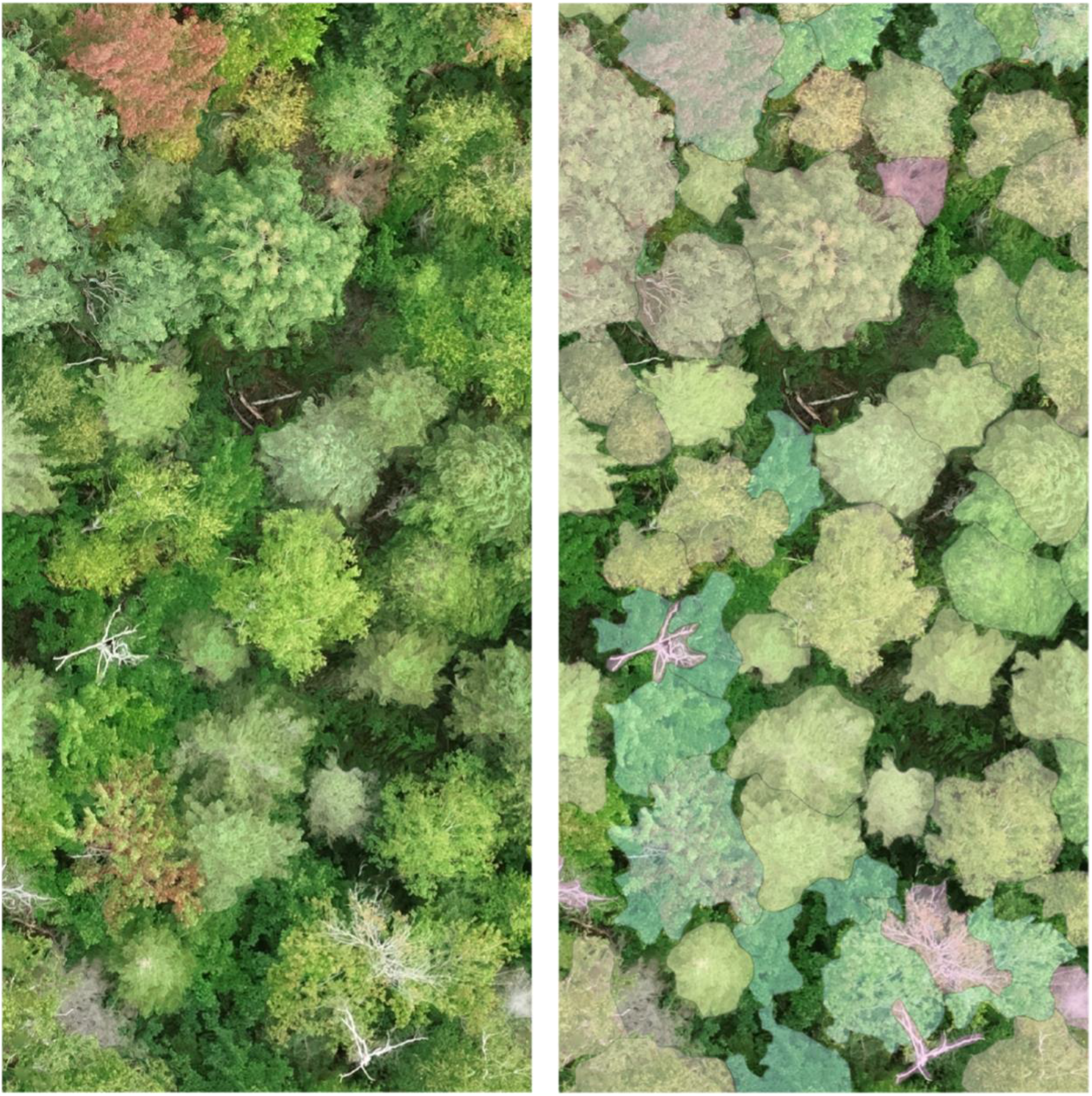
Image of the annotations from the dataset. On the left is a section of the orthomosaic representing the canopy of the forest on the 2^nd^ of September. On the right are the annotations segmenting the tree crowns on the orthomosaic.

The polygons delineating the tree crowns were only drawn using the orthomosaic that was the clearest (i.e diffuse light conditions, minimal artefacts, leaves still on), which was the imagery taken on September 2^nd^, 2021. We decided to create a single layer of polygons instead of one layer per imagery due to the very long time it took to complete the annotations. Since the orthomosaics were very accurately georeferenced (generally <3 cm error), the polygons could be applied to the other dates with minimal shifts (Figure S2). We did not adjust the polygons for the different dates to account for fallen trees or other major changes as we considered such changes to be relatively rare and would thus not affect the overall quality of our dataset.

The classes used are presented in Figure 3. These 14 classes represent the crowns that were the most abundant at the top of the canopy. The proportion of the area covered by each species for the training and test sites is shown in Table 1. Most of the classes (11) are at the tree species level, two of them are at the genus level and there is also one class for dead trees. The dataset includes six classes representing conifers and seven classes representing deciduous broadleaf species. In the analyses and the discussion, *Larix laricina* (Du Roi) K. Koch, a deciduous conifer, has been included with the evergreen conifers as the imagery stops before the senescence starts for this species.

**Figure 3.**
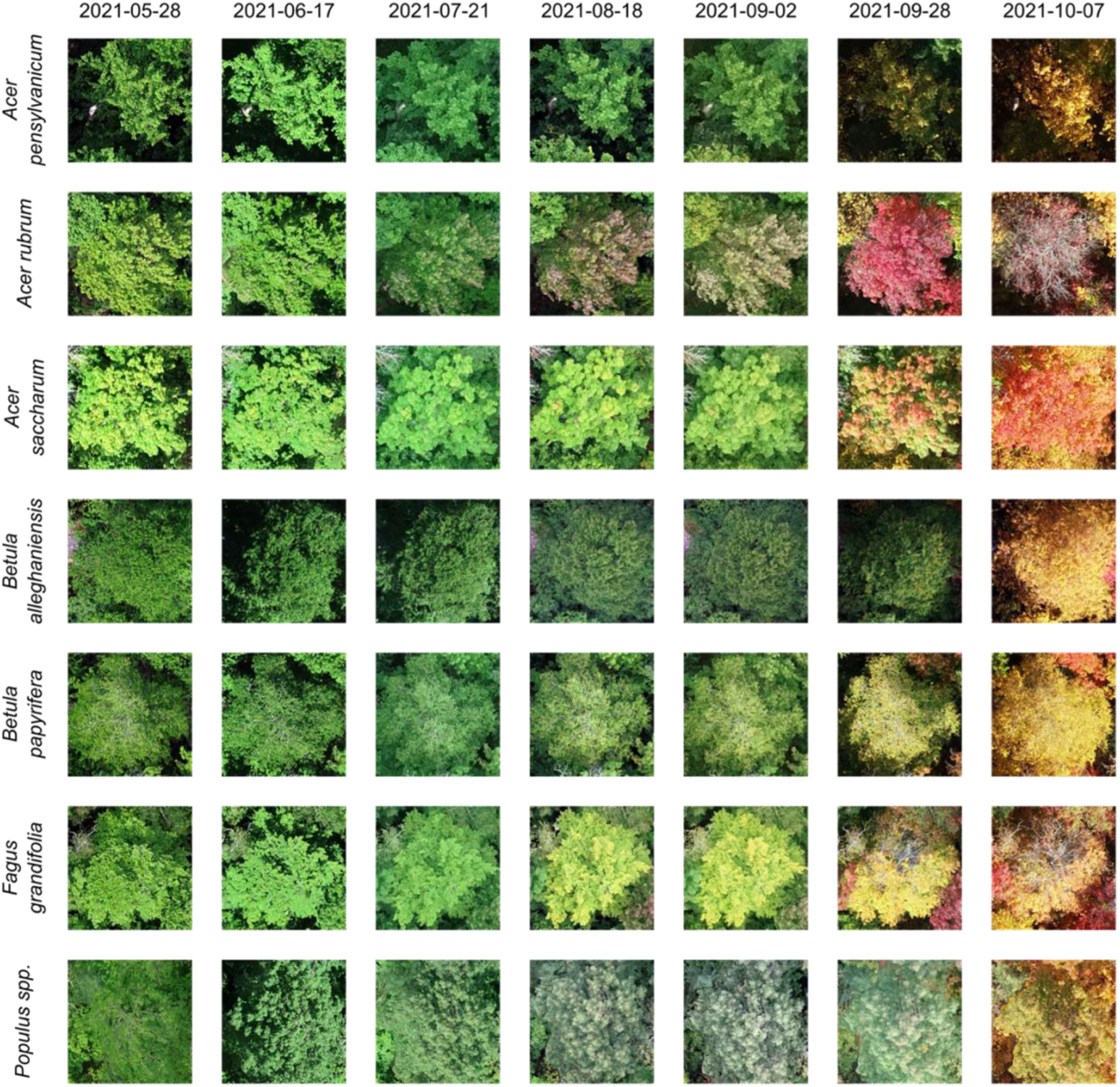
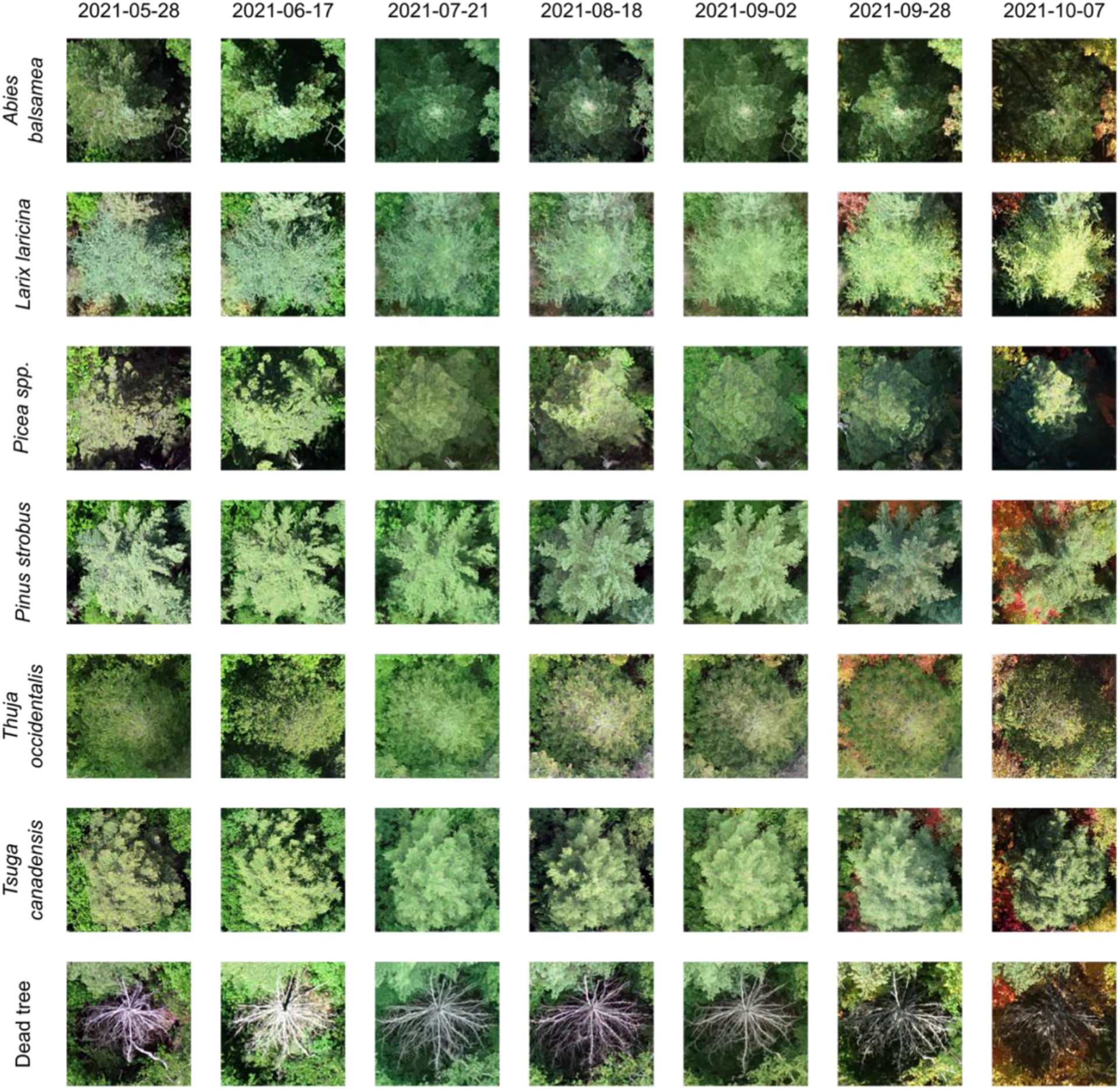
Example of the fourteen classes from the seven orthomosaics (May to October) The first half represents the seven broadleaf species, and the second half represents the six conifers and the dead trees. The same crown is shown for each species from May to October.

**Table 1.** F1-score for the models for each class at each date. This represents the performance of the trained model after applying it to new independent imagery (test dataset). The highest F1-score for each date is in bold. The highest average F1-score for the dates and classes is also in bold.

| F1-score | Proportion of<br>pixels (%) | Date |  |  |  |  |  |  | Average |
| --- | --- | --- | --- | --- | --- | --- | --- | --- | --- |
|  |  | 2021-05-28 | 2021-06-17 | 2021-07-21 | 2021-08-18 | 2021-09-02 | 2021-09-28 | 2021-10-07 | F1-score |
| *Abies balsamea* | 5.1 | 0.69 | 0.68 | 0.64 | 0.69 | 0.74 | 0.66 | 0.63 | 0.68 |
| *Acer pensylvanicum* | 1.8 | 0.61 | 0.55 | 0.52 | 0.41 | 0.54 | 0.45 | 0.36 | 0.49 |
| *Acer rubrum* | 23.1 | 0.61 | 0.61 | 0.57 | 0.62 | 0.64 | 0.63 | 0.55 | 0.60 |
| *Acer saccharum* | 7.2 | 0.60 | 0.59 | 0.57 | 0.61 | 0.60 | 0.64 | 0.54 | 0.59 |
| *Betula alleghaniensis* | 2.6 | 0.53 | 0.65 | 0.62 | 0.69 | 0.74 | 0.63 | 0.66 | 0.65 |
| *Betula papyrifera* | 31.1 | 0.69 | 0.70 | 0.68 | 0.71 | 0.79 | 0.73 | 0.72 | 0.72 |
| *Fagus grandifolia* | 1.3 | 0.59 | 0.52 | 0.56 | 0.60 | 0.71 | 0.43 | 0.39 | 0.54 |
| *Larix laricina* | 0.5 | **0.80** | **0.81** | 0.75 | **0.80** | 0.79 | 0.73 | 0.75 | 0.77 |
| *Picea spp.* | 2.2 | 0.69 | 0.67 | 0.61 | 0.67 | 0.73 | 0.69 | 0.65 | 0.67 |
| *Pinus strobus* | 6.3 | 0.76 | 0.79 | **0.79** | 0.75 | **0.87** | **0.77** | **0.77** | **0.79** |
| *Populus spp.* | 13.5 | 0.72 | 0.76 | 0.72 | 0.75 | 0.83 | **0.77** | **0.77** | 0.76 |
| *Thuja occidentalis* | 3.2 | 0.66 | 0.66 | 0.65 | 0.66 | 0.72 | 0.68 | 0.64 | 0.67 |
| *Tsuga canadensis* | 0.3 | 0.65 | 0.63 | 0.61 | 0.66 | 0.75 | 0.61 | 0.53 | 0.63 |
| Dead tree | 1.6 | 0.54 | 0.57 | 0.54 | 0.61 | 0.62 | 0.54 | 0.52 | 0.56 |
| Average F1-score |  | 0.65 | 0.65 | 0.63 | 0.66 | **0.72** | 0.64 | 0.61 |  |

#### 2.2.3 Data splitting and tiling

To train a semantic segmentation model, the orthomosaics and the polygons need to be separated into regular tiles and masks. The tiles correspond to a section of the orthomosaic and the mask to the related annotations. We ran the tiling of the orthomosaics and the annotations in R v.4.1.1 (R Core Team, 2021) where we created a grid covering the entire area and selected tiles where at least 25% of the surface was covered by annotations. This avoided including in the dataset regions where the orthomosaic was not clear enough to annotate. To train our models, we chose non-overlapping tiles of 256 x 256 pixels based on the results from Schiefer et al. (2020), which corresponds to approximately 5 x 5 m on the ground. Examples of such tiles and associated masks are shown in Figure S3.

The tiles in the training site were randomly divided between training data (80%) and validation data (20%). The tiles in the test dataset (approximately 20% of the entire dataset) contained the annotations from the test site as well as some annotations from the training site (which were excluded from the training data) to balance the classes (Figure 1). This was done because some of the less abundant species were all present in the training site, but not in the test site. The dataset used will be available soon at https://doi.org/10.5281/zenodo.8148479.

### 2.3 Deep learning model and training

The type of model used for training is a CNN, specifically a U-Net (Ronneberger et al., 2015), which is a semantic segmentation algorithm. We chose U-Net as our architecture because of its success in other individual tree classification problems (James & Bradshaw, 2020; Korznikov et al., 2021; Schiefer et al., 2020; Wagner et al., 2019). We trained one model for each date and evaluated the performance of this model on the test data from the same date. We did not apply models trained on one date to other dates as this was out of the scope of the study. The parameters used for training were the same for all models {batch: 14, learning rate: 0.001, epochs: 60, optimizer: Adam}. We utilized data augmentation techniques to enhance the robustness of our dataset and to account for the high heterogeneity. Specifically, we applied random rotations with a probability of 0.5, along with horizontal and vertical flips with a probability of 0.75, directly on the batch dataset.

All the code for the training and the testing was written in Python v.3.9 (Python Core Team) using the fast.ai library (Howard & Gugger, 2020a). The fast.ai library is a high-level library for deep learning models based on PyTorch. While it limited us regarding certain parameters, it allowed us to train the models using a more accessible workflow (Howard & Gugger, 2020b). We ran the code on Google Colab Pro+ which gave us access to NVIDIA A100 Tensor Core GPUs.

### 2.4 Accuracy assessment

To assess the performance of the models, we used the F1-score. This metric is calculated by taking the harmonic mean of the precision and recall (Equation 1). The recall represents the proportion of actual presences that were correctly classified, and the precision represents the proportion of predicted presences that were classified correctly. When applied to the test dataset, we extracted the F1-score for each of the classes and calculated the average per date and per class.

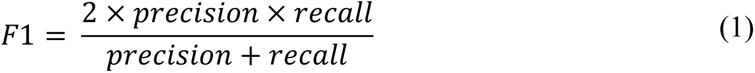

We then represented the accuracy of the predictions using confusion matrices by calculating the overlap between the annotations and the predictions from the model and calculating the percentage. The confusion matrices allowed us to see which pixels were misclassified and between which classes for the different dates. By comparing the confusion matrices and the F1-scores, we could determine how the phenology of each species during the growing season can affect the performance of the models.

## 3. Results

### 3.1 Model results

The model that showed the highest overall average F1-score (0.72) was trained on imagery from early September. The second-best model was trained with imagery from August, with an F1-score of 0.66. The other models showed an average F1-score between 0.65 and 0.61. The model trained on the October imagery had the lowest F1-score, with 0.61.

The classes with the highest F1-scores are *L. laricina*, *P. strobus*, and *Populus* spp. *Pinus strobus* had the highest average F1-score for all the dates. The F1-scores for *P. strobus* range between 0.77 and 0.87, where the lowest scores were from the models trained with the imagery taken in late September and October. The classes with the lowest F1-scores were *A. pensylvanicum*, *F. grandifolia* and *Betula alleghaniensis*. *Acer pensylvanicum* had the lowest overall average F1-score for all dates. The F1-scores for *A. pensylvanicum* ranged between 0.36 and 0.61.

When comparing the deciduous broadleaf species and the evergreen species (including *L. laricina*, which we grouped with the evergreen trees), general trends regarding the variation of the F1-score throughout the growing season can be seen between the classes from each group (Table 1, Figure S4). The evergreen trees showed less variability in their F1-score through time, and they all showed similar trends, except for *P. strobus* and *L. laricina* which had higher F1-scores overall. *Tsuga canadensis* had a lower score for the imagery from late September and October. As for the deciduous broadleaf classes, *Populus* spp. and *B. papyrifera*, the first and second best predicted deciduous classes, have trends very similar to the ones seen in the evergreen classes. *Acer pensylvanicum* and *F. grandifolia*, the two worst predicted deciduous classes, along with *B. alleghaniensis* have more variability in their score and different trends from any other class. *Betula alleghaniensis* is the only deciduous class with an increase in its F1-score in October. Three of the deciduous classes, *Populus* spp, *B. papyrifera* and *A. rubrum*, have higher scores in September (early and late) compared to the rest of the dates. *Acer pensylvanicum*, *A. rubrum*, *A. saccharum* and *F. grandifolia* showed a decline in their F1-score from the predictions done on the imagery from late September and October.

### 3.2 Predictions errors in confusion matrices

The diagonal in the confusion matrices (Figure 4) corresponds to the true positive for each class. These values were high, meaning the label for most of the predicted pixels corresponds to the label of the annotated pixels. This is the case for the matrices comparing the predictions to the annotations for each of the seven dates (Figure S5).

**Figure 4.**
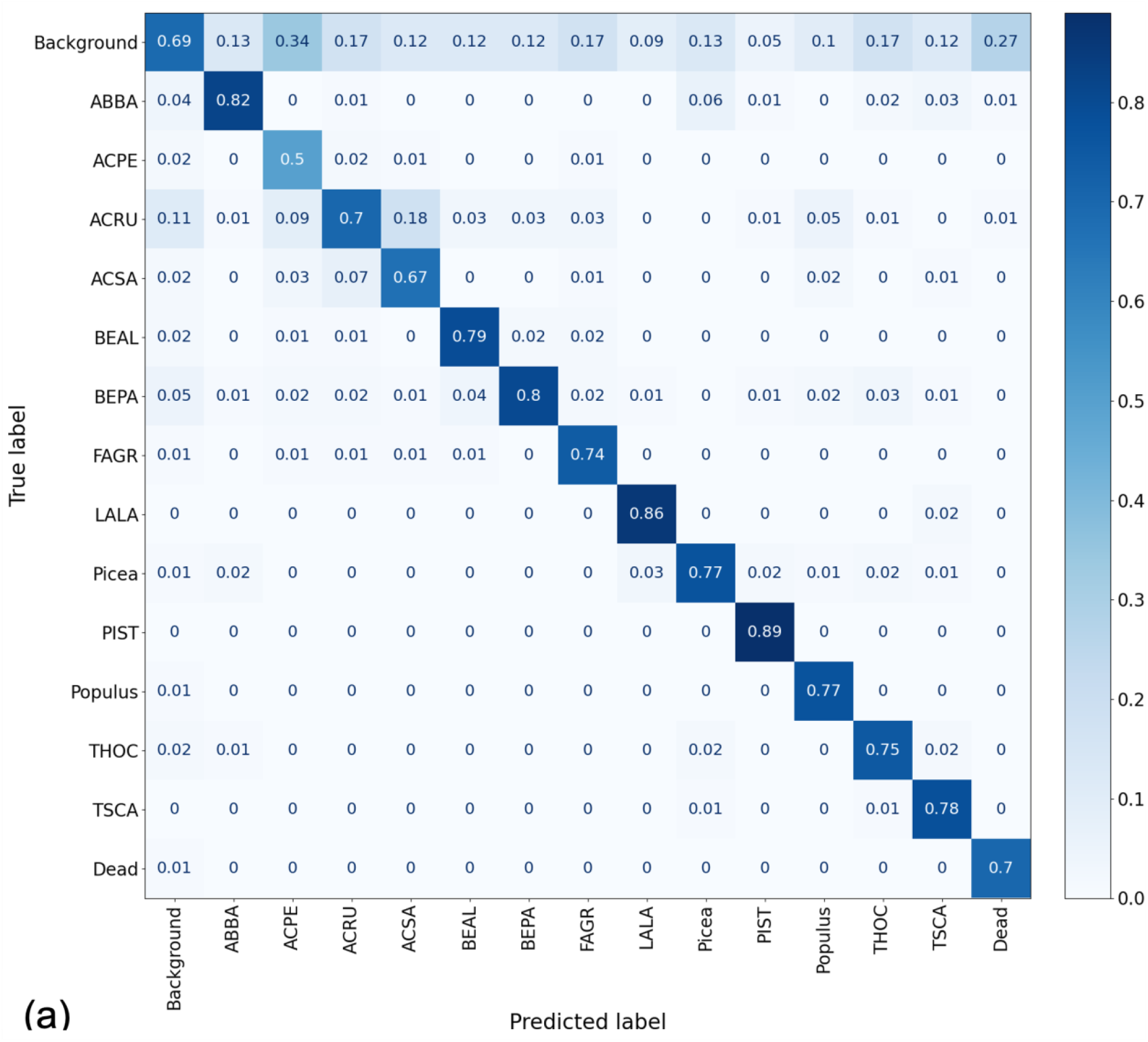
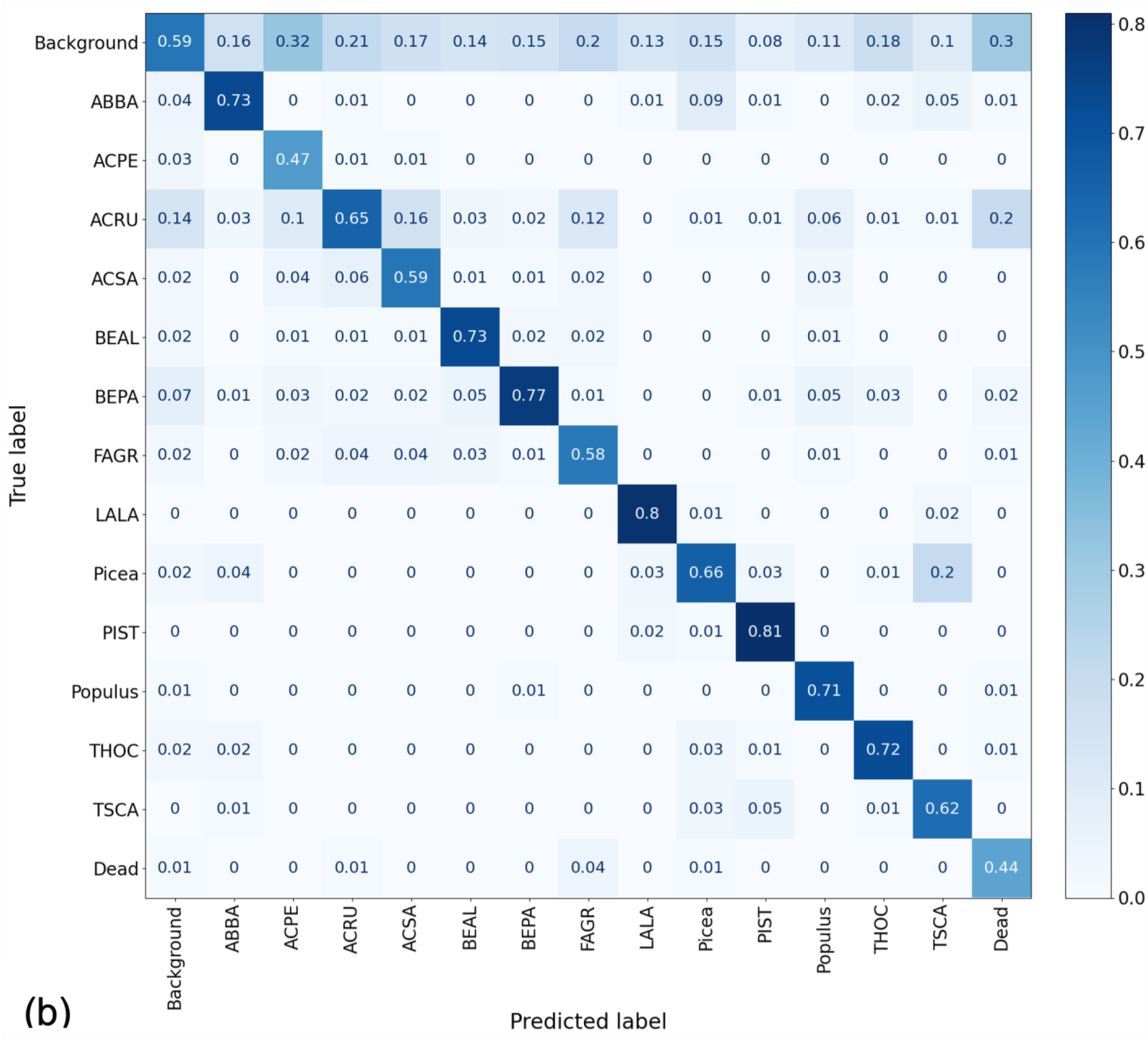
Confusion matrices for early September and October. The values represent the overlap (from 0-1) between the annotations (true label) and the predictions (predicted label) done by the model on unseen imagery from (a) September 2^nd^, 2021, and (b) October 10^th^, 2021. ABBA: *Abies balsamea*, ACPE: *Acer pensylvanicum*, ACRU: *Acer rubrum*, ACSA: *Acer saccharum*, BEAL: *Betula alleghaniensis*, BEPA: *Betula papyrifera*, FAGR: *Fagus grandifolia*, LALA: *Larix laricina*, Picea: *Picea spp.*, PIST: *Pinus strobus*, Populus: *Populus spp.*, THOC: *Thuja occidentalis*, TSCA: *Tsuga canadensis*.

The classes with the highest performances in the matrices were mostly consistent with what was seen with the F1-score in Table 1. *Pinus strobus* (Figure 5a) and *L. laricina* (Figure 5b) had the highest values in the matrices overall, followed by *B. papyrifera* and *A. balsamea*. On the other hand, *A. pensylvanicum*, dead trees and *F. grandifolia* were the classes with the lowest values in the matrices overall.

**Figure 5.**
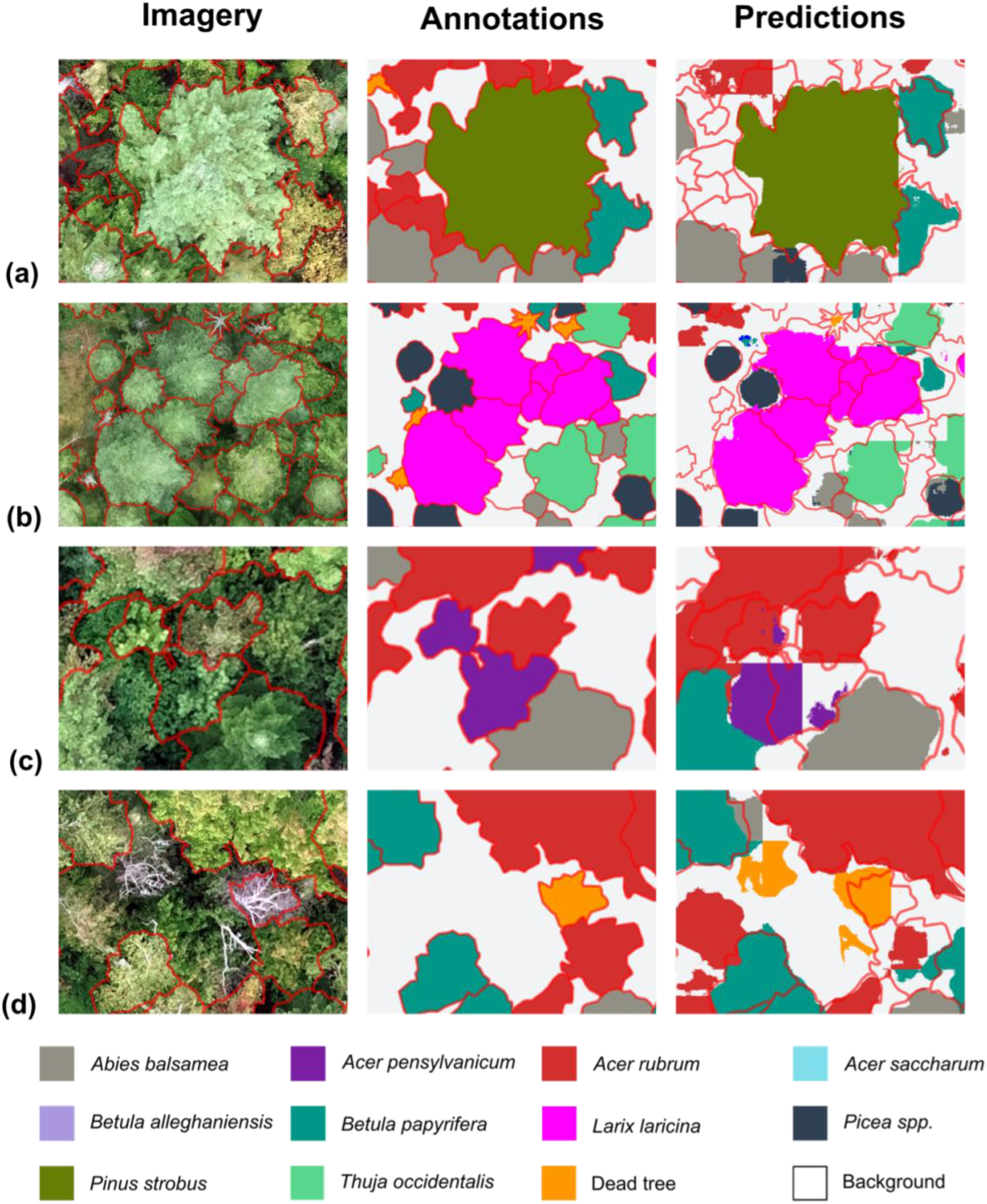
Classes with higher and lower performances in the predictions. The classes with the highest performance in the predictions were (a) *Pinus strobus* and (b) *Larix laricina*. Two of the classes with the lowest performance in the predictions were (c) *Acer pensylvanicum* and (d) dead trees. These predictions were generated by the model trained with the imagery from September 2^nd^, 2021.

In general, the models overestimated the number of pixels for each class, in that predictions for each class included what was labelled as background. This is illustrated in the confusion matrices by larger values in the top row. This overestimation of the number of pixels representing each class can be seen throughout the season.

Some classes had consistently lower values of pixels misclassified with background, such as *P. strobus*. Other classes had consistently higher proportions (> 27%), mainly *A. pensylvanicum* and dead trees. As mentioned earlier, *A. pensylvanicum* showed the lowest performance of all the classes. The predictions were less accurate when crowns are surrounded by other vegetation than when the crown is more isolated and distinct from its surroundings (Figure 5c). Dead trees also had a low performance when looking at the values in the confusion matrices. The imagery contained a lot of dead branches along with full crowns and many of the predictions for this class included these single, isolated branches as well as partially dead crowns (Figure 5d).

The main classes that were misclassified were the maple species, *A. pensylvanicum*, *A. rubrum* and *A. saccharum*. Specifically, pixels labeled as *A. rubrum* in the annotations were often predicted as being *A. saccharum* (Figure 6a) and, to a lesser extent, *A. pensylvanicum*. This difference in the annotations and the predictions is present in similar proportions for all the dates, with only the model trained with the July imagery having a higher proportion of misclassifications between the maples. In a smaller proportion than for the maples, the two birch species were also misclassified where *B. papyrifera* pixels are being predicted as *B. alleghaniensis*. In some cases, classes from different genera were confused by the models as well. *Acer rubrum* was mistaken for *F. grandifolia* for all the dates, but especially in the early-mid part of the summer (June and July) and in the fall (end of September and October). From May to early September, a very small proportion (1-2%) of pixels labelled as *A. rubrum* were predicted as being dead trees. This proportion increased to 10% for the end of September and 20% for October (Figure 6b).

**Figure 6.**
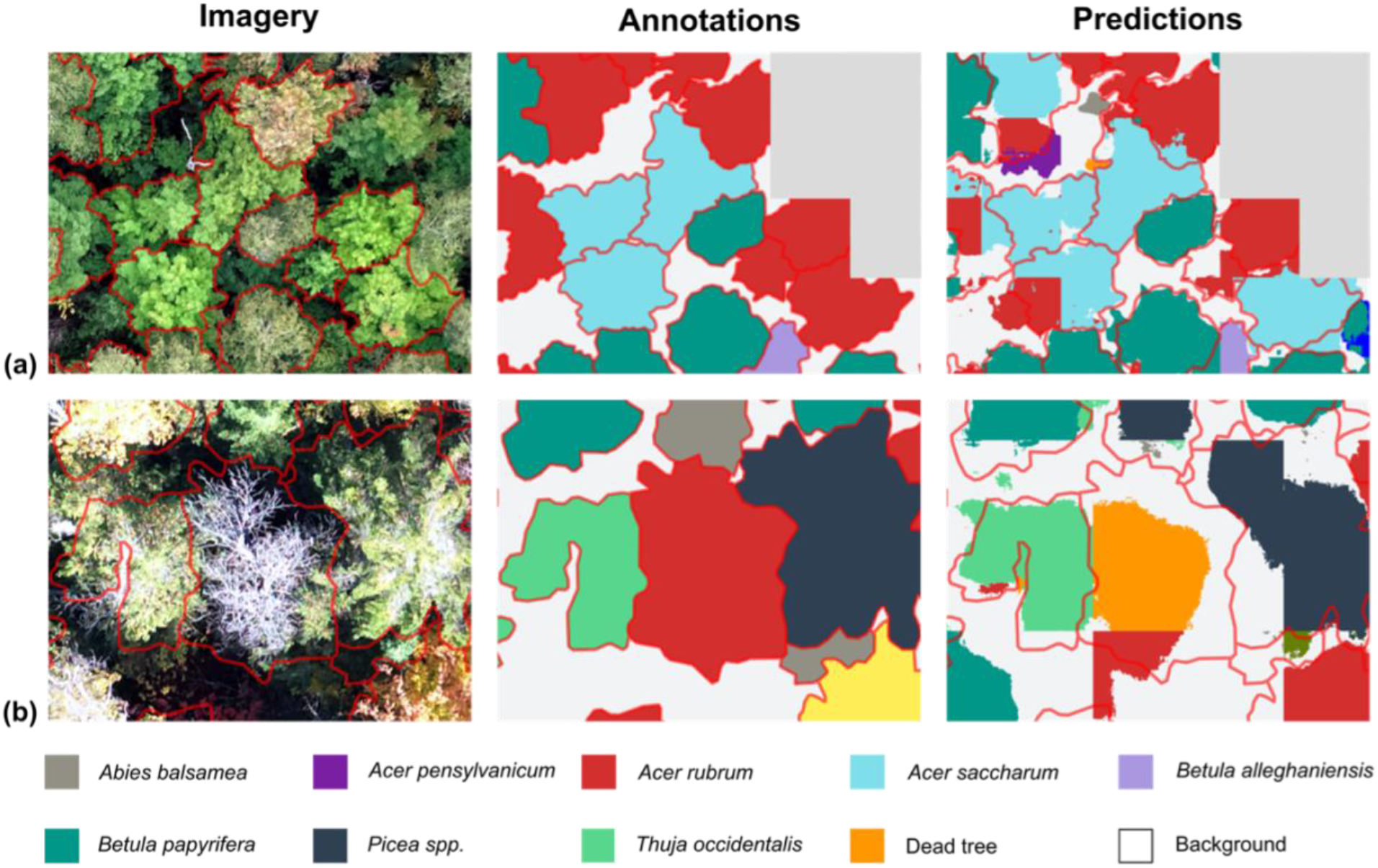
Classes that showed the highest proportions of misclassifications. Some classes had high proportions of misclassification with a class other than the background such as (a) the maples, specifically *Acer rubrum* and *Acer saccharum* all throughout the season (example from 2^nd^ of September), and (b) *Acer rubrum* often identified as a dead tree in October. The parts in grey in (a) were not included in the training or the testing of the model as the annotations covered less than 25% of the area of those tiles.

## 4. Discussion

Our results showed that semantic segmentation on UAV imagery can accurately classify tree species in a temperate forest, as was previously highlighted in other studies (James & Bradshaw, 2020; Kattenborn et al., 2019; Schiefer et al., 2020). However, our results offer new insights about the temporal variation in tree species classification driven by leaf phenological changes. Contrary to our hypothesis, tree species classification was not more accurate during peak autumn colours (i.e. early October) in this temperate forest. Instead, classification accuracy was highest at the start of the senescence process in early autumn, when autumn foliage colours started to appear (i.e. early September), and was in fact lowest for most species at peak autumn colours in early October. Still, classification accuracy remained relatively high throughout the growing season, even in early summer where tree foliage appeared more uniformly green, among and within tree species. Together, our results provide further evidence of the power of high-resolution UAV imagery and deep learning to classify tree species in temperate forests, but shows how the development of leaf colour during senescence can influence the performance of such classification models.

### 4.1 Lowest accuracy for October imagery

The model trained and tested with the October imagery showed the lowest F1-score out of all the dates. This went against our initial hypothesis, since we expected that imagery at peak autumn leaf colours would give the best predictions. Our results differ from those of previous studies seeking to determine the optimal timing for aerial imagery to differentiate tree species, which concluded that imagery acquired in autumn, when species display the largest colour differences, gave the highest accuracy (Grybas & Congalton, 2021; Hill et al., 2010; Key et al., 2001; Weil et al., 2017). However, in the early and late September models, when leaf colouring had started but was not at its peak, the F1-scores were higher than for the October model. This suggests other aspects of senescence beyond colour change, such as leaf abscission and drop, can also influence tree species classification from UAV imagery. On the imagery from October, it was possible to see the forest floor and vegetation below the canopy through the tree crowns that had lost most of their leaves. This can affect the appearance of the imagery and increase the influence of the background (forest floor, sub-canopy vegetation) within the imagery (Cho et al., 2012). It could also change the crown structure by exposing primary and lateral branches on the imagery (Veras et al., 2022). Together, our results suggest that peak autumn foliage colour does not lead to the highest tree species classification accuracy from UAV imagery in a temperate forest. Instead, the highest performance was obtained at the start of the senescence process when leaves were just starting to change colour.

We attribute a large part of the drop in accuracy for the October model to the leaf fall that happens during senescence. Deciduous tree species classification accuracy decreased from September to October, particularly for tree species that had then lost most of their leaves, such as *A. rubrum* and *F. grandifolia*. Similarly, this was observed in the Amazonian forest during the dry season, where there was a drop in the CNN accuracy relative to other phenological periods (Veras et al., 2022). The authors associated this decrease in performance with significant leaf drop during the dry season. However, it is important to note that in that Amazonian study as well as ours, species classification accuracies remain high for most deciduous tree species, despite such variation in crown appearance caused by leaf drop.

While some authors have recommended to avoid acquiring imagery when most of the leaves have fallen (Wolter et al., 1995), recent studies using CNNs to map tree crowns suggest that including more leafless trees during training could help the model to generalize to images taken throughout the year (Braga et al., 2020). For instance, our October model seems to have been able to learn features of leaf-on and leaf-off crowns of *A. rubrum*, one of the most abundant species in our dataset and one of the species with significant leaf fall in the October imagery. Some *A. rubrum* leafless crowns were correctly identified at least in part, but more leafless crowns would be necessary to adequately generalize the features for the CNN to identify both leaf-on and leafless crowns. Another species with significant leaf fall in October was *F. grandifolia*. However, because of its low abundance in the dataset, many *F. grandifolia* crowns have not been detected or were misclassified with the background or other deciduous species. To avoid a decrease in prediction accuracy in autumn, more leafless crowns may be needed to train the model to learn species-specific branching patterns. Additionally, the branching structure may have less distinguishable features due to less contrast with the surroundings making it more difficult to distinguish from the background, and as such may require more training data for the model to correctly classify them (Cho et al., 2012; Kattenborn et al., 2020). So, while senescence and leaf colouring in autumn in temperate forest can add contrast in terms of colour among tree species and can be a useful trait to differentiate them (Fassnacht et al., 2016), the timing of leaf fall also differs within species. Intraspecific variability in the timing of leaf fall can lead to a decrease in the accuracy of the model since it increases within-class heterogeneity.

### 4.2 Challenges related to the variability of senescence

Our initial hypothesis was that tree species classification accuracy from UAV imagery would be highest at peak autumn colours when the contrast in foliage colour among species was expected to be highest. We defined the 7^th^ of October as the peak date for colouring from our field observations and based on previous phenocam observations at that site (see Laurentides site at http://phenocam.unh.edu). However, this decision was subjective and based on visual interpretation of the level of colouring and leaf fall in the overall landscape, while there can be significant spatial and temporal variability in senescence even within landscapes.

The definition of senescence is broad, ranging from the start of leaf colour change to leaf abscission, which includes multiple events that should not be assigned to a single date and that should be treated as a multiple-day event (Denny et al., 2014; Gallinat et al., 2015). Due to differences in species composition and environmental conditions in different studies, it is difficult to compare the timing and the dates chosen to acquire the imagery. In fact, the beginning of leaf fall varies between species (Veras et al., 2022), which we observed in the target species. This can be due to environmental conditions, especially considering the variable topography and the variety of soil types and conditions in our study site (Courchesne & Hendershot, 1989; Savage, 2001). While we did not measure soil conditions, these topo-edaphic conditions could have influenced the timing of senescence. Temperature variations associated with microclimate from the topography can also affect phenology on smaller scales (Lechowicz, 1984; Lisein et al., 2015; Richardson & O’Keefe, 2009). Moreover, abundant soil moisture can delay leaf colouring and leaf drop while drought and low moisture levels could advance senescence (Gallinat et al., 2015; Leuzinger et al., 2005). Early frost and high winds can also advance tree senescence depending on topographical position (Chmielewski et al., 2005; Norby et al., 2003). This combination of conditions affect the trees and their leaves throughout the growing season, causing leaf senescence to be an asynchronous event between and within species in heterogeneous areas (Gallinat et al., 2015). Therefore, classifying deciduous species in a temperate forest during senescence can be made difficult due to challenges related to predicting the timing of leaf colouring and leaf fall (Onishi et al., 2022).

Some tree species, such as *A. rubrum*, can tolerate a wide range of environmental conditions (Abrams, 1994; Fowells et al., 1965), which might influence its phenological response in autumn (Klosterman et al., 2018). The occurrence of *A. rubrum* in a wide range of conditions may explain the decline in the F1-score of that species in October. This could have led to different timing for the colouring and falling of leaves depending on the specific environmental conditions, causing more variability in the appearances of crowns. Indeed, an important factor that can negatively affect tree species classification performance is the colour variation of the leaves within a species, which tends to be greater in mixed forests (Lisein et al., 2015). Therefore, in a region with a variable topography and a diverse tree species composition such as ours, the timing of phenological events may vary significantly both among and within species in response to these conditions, which can influence tree species accuracy from UAV imagery.

Compared to autumn phenology and leaf senescence, spring leaf phenology is more homogeneous in terms of colour response than autumn leaf phenology. Indeed, spring phenological events tend to be more synchronized within species and the different species are distinct enough to differentiate between them (Lisein et al., 2015). We acquired imagery at the end of May, during late leaf flushing, and the accuracy of that spring model was higher than those from late September and October. Tree species such as *A. rubrum* and *F. grandifolia* were better classified in May than in October, pointing to higher homogeneity in spring leaf phenology for these species compared to autumn phenology. However, the opposite pattern was found for species where leaf colouring happened later in the season, such as *B. alleghaniensis*, which had better predictions in September and October than in May. While the objective of this study was not to determine an ideal date for imagery to map autumn phenology, our results highlight the fact that because of the many factors influencing the timing of senescence, especially regarding leaf fall, a single date cannot be chosen to capture peak colouring for multiple species in a mixed heterogenous forest. The use of multitemporal imagery with higher temporal resolution (e.g. every few days) during senescence would take advantage of the capacity deep learning models have to learn generalized features from imagery and make the models more adapted to variability within the imagery (Natesan et al., 2020), such as the timing of senescence or light conditions.

### 4.3 Tree segmentation in a complex canopy

Field observations combined with visual interpretation of the orthomosaic are necessary to build an extensive, accurate and precise segmentation dataset in a complex forest canopy to train CNN models (Kattenborn et al., 2021). In our study, having visual interpreters allowed us to segment tree crowns with irregular shapes, others that were partially covered by other crowns or that were overlapping other trees from the same or different species. Other studies that used visual interpretation to build a dataset of tree crowns also highlighted the value of this approach, even if there can be some misinterpretation of the imagery (Beloiu et al., 2023; Schiefer et al., 2020). Additionally, when annotating the imagery directly with this method, it ensures an accurate georeferencing of the annotations aligned with the imagery (Schiefer et al., 2020), which is especially important when working with multitemporal imagery.

The quality of the labelling and the delineation of the tree crowns has a direct influence on the quality of the species classification (Fassnacht et al., 2016). Some CNNs have been able to compensate for mislabelled crowns by correcting the classes in the predictions, but only to some extent (Hamdi et al., 2019; Kattenborn et al., 2020). Moreover, these corrections would also affect the metric of the model as they would not correspond to the training data. Therefore, it is best to avoid faulty labels and inaccurate segmentations, but certain characteristics of the target species can lead to mistakes. For many species, including most broadleaf species and some conifer species such as *T. occidentalis*, the borders of the crowns are not clearly identifiable on the imagery which may affect the accuracy of the annotations and the predictions (Gan et al., 2023). While delineating the crowns, the border of the crowns may vary slightly depending on whether this is done by a visual interpreter or the CNN, leading to differences in the labelling of the pixels on the borders of the crowns. Additionally, closed and overlapping canopies are more difficult for delineating tree crowns than for isolated trees or uniformly and distributed planted trees (Nevalainen et al., 2017; Yang et al., 2022). Tree detection algorithms are also known to be less accurate in forest with complex crown structures (Nuijten et al., 2019) or heterogenous conditions such as a higher diversity of species and shapes (Berra, 2020; Hastings et al., 2020), which is the case for most of our study site and makes the segmentation of tree crowns a laborious task.

We chose to annotate only one of our multiple dates, an approach that was also followed by Veras et al. (2022). This was done to reduce the time required to annotate all the imagery while still ensuring that the annotations were of high quality by ensuring that annotations and imagery were aligned for all dates (Figure S2). The cm-level alignment of all imagery among dates was made possible by using a UAV equipped with an RTK GNSS receiver in combination with GCPs. It is unclear whether the use of a single date for our annotations might explain the higher performance of the early September imagery or if other attributes of the imagery, such as light conditions or the phenology of the species, had a stronger influence on the performance. While additional data would be needed to test for this, we can see that the proportion of background pixels that have been classified as each target class, corresponding to the false positives, is relatively high for all the dates. This leads us to believe the models are less accurate in predicting the pixels on the border of the polygons. The accuracy of the predictions can be affected by the edges of the tree crowns which are often not well defined or clear (Gan et al., 2023). This could also be because the models are correcting for some missing labels in the imagery. On the other hand, the models made relatively few mistakes between the classes. If the predictions from the early September model would have been influenced by the alignment of the annotation, we would expect a significant decrease in false positives, which was not the case. More tests would be necessary to explore this, but we believe that the higher F1-score from early September is not only attributed to the fact that the annotations were made from this date.

Recent computer vision foundational algorithms such as the Segment Anything Model (SAM, Kirillov et al., 2023) may help to reduce the time necessary to build such extensive segmentation training datasets. However, it is still unknown how well such algorithms would perform on complex forest canopies, with overlapping crowns and fuzzy contours. In medical imagery, the performance of SAM was tested and gave variable results depending on the imagery and the parameters used, which led to the conclusion that some fine-tuning may be necessary for different applications (Huang et al., 2023). More research would need to be done to assess the capabilities of SAM in this context and to determine if this model were a promising tool in remote sensing of forest canopies.

The dataset created here (23,000 segmented individual tree crowns) includes 13 species and genera, which have an environmental and an economic importance in Northeastern North America. Moreover, these trees are segmented on seven different dates, totalling almost 161,000 annotated tree crowns. Due to the nature of the annotations, other deep learning approaches may be used such as instance segmentation, which detects the individual object segments them (Kattenborn et al., 2021). This dataset is available online for use by others. The workflow presented here could be applied to other types of forests to assess the effects of the phenology specific to the region. With recent advances in segmentation tools, such as SAM (Kirillov et al., 2023), building a dataset for this type of method may become increasingly accessible depending on the complexity of the forest and the capacity of these types of models. More high temporal resolution segmented imagery of forest canopies can significantly improve our ability to accurately map and monitor these ecosystems.

### 4.4 Other features from the imagery

Different characteristics of the imagery, such as illumination conditions, spatial resolution, and texture, can influence the performance of the model (Flood et al., 2019; Kattenborn et al., 2019). Light conditions can influence the quality of the predictions of the CNNs, especially when comparing the F1-score for dates with diffuse light conditions (e.g., early September and August) and dates with full sun conditions (e.g., July and October). This may be combined with other features of the imagery, such as the effects of phenology discussed earlier. Since we did not test directly for the effect of different illumination conditions, we cannot say whether the light conditions or the phenology had a stronger impact on the F1-score. Some authors have stated that the changing tree phenology has as a greater impact on the quality of their imagery than illumination conditions (Lisein et al., 2015). Other studies using a CNN to detect tree crowns have reported a minor influence of illumination conditions on the classification (Beloiu et al., 2023), while others have reported an important decrease in accuracy due to shaded areas in the imagery when using a maximum-entropy classifier (Lopatin et al., 2019). This seems to indicate that CNNs are more robust to changes in illumination conditions and are still able to extract the relevant information, despite shaded areas. If the crown is distinguishable on the imagery, meaning it is not completely dark, the model is likely to identify it since the conditions were the same in the datasets used for training and testing. In fact, deep learning algorithms may be better at coping with shadows in the imagery, using them as an additional source of species-specific information, especially regarding the structure of the canopy (Kattenborn et al., 2020; Lopatin et al., 2019). Despite this, when possible, it might be best to avoid strong cast shadows in the imagery, especially in a forest with trees of variable heights, which could exacerbate the effects of shadows and make the segmentation task more difficult.

The spatial resolution of the imagery can influence the model’s ability to classify tree species. When a CNN is trained with high-resolution imagery, it increases the accuracy as it can extract more contextual details in the imagery such as leaf form or branching patterns, which may be necessary to differentiate between species (Kattenborn et al., 2019). Higher spatial resolution also increases the contrast in texture that is visible on the imagery (Park et al., 2019). Low-contrast regions could lead to less accurate predictions from a CNN, showing the importance of texture in the imagery (Flood et al., 2019). When comparing the importance of texture and colour for a classification problem, the model that used texture features outperformed the model that was based only on colour. It was concluded that texture should be used when colour features do not perform well enough since they can be more robust to different light conditions than colour (Park et al., 2019).

In high-resolution imagery, for a given pixel in a tree crown, its neighbouring pixels are likely to be in the same crown and have similar information and therefore be classified as the same class (Woodcock & Strahler, 1987). However, this was not always the case. An edge effect, where the edges of the tiles result in artefacts in the predictions, can often appear when the CNN cannot determine with high confidence the label of a crown. This results in some crowns being assigned different classes depending on the tile, affecting the accuracy of the predictions (Flood et al., 2019). This was especially true for classes that were close phylogenetically with similar features such as *A. rubrum* and *A. saccharum*. For these species, some crowns were identified as both classes, often separated by a line representing the tile edge. While higher spatial resolution brings more information to the imagery, it also increases within-class variability due to increased spatial detail, canopy structure and shadows (Hay et al., 1996; Lopatin et al., 2019). This could lead to misclassifications especially in classes with overlapping features. Finally, higher-resolution imagery also entails higher computational costs.

### 4.5 Species-specific traits

Distinctive traits as well as constant traits within a class can play an important role in the performance of a tree species classification model. Some of these traits may be associated with the species, the functional traits or type, in particular deciduous broadleaf tree species and conifers. While autumn leaf colouring can help to distinguish between deciduous trees and evergreen trees (Onishi et al., 2022), our results show that deciduous broadleaf species consistently show a lower average F1-score than conifer species, regardless of the date. Other studies have also reported a higher performance for conifers than broadleaf species for segmentation and tree counting (Onishi et al., 2022; Zhang et al., 2022). The difference in crown structure between broadleaf species and conifers and the variability from one individual to the other could explain this. Deciduous trees tend to have a more complex and plastic crown structure, while conifers have a more conical crown and a stronger relationship between height and crown size, leading to more of a constant crown structure (Nuijten et al., 2019). This was the case with the conifers in our dataset, which did not vary as much as broadleaf trees during senescence. As such, conifer species with homogeneous characteristics throughout the growing season are more likely to be better detected by the model (Veras et al., 2022), and they are not affected by the asynchronous timing of senescence from one individual to another. Two broadleaf classes stood out from the rest in that they showed a higher F1-score, *Populus spp.* and *B. papyrifera*. Both tree species are intolerant to shade and are therefore always at the top of the canopy with their crowns completely visible and not affected by the shade of neighbouring trees. *Populus spp.* has more constant features from one crown to another, while *B. papyrifera* showed more variability in crown shape and timing of leaf colouring and drop which may have been compensated by the large number of annotations for this species.

Some classes may be easier to classify by the model due to distinctive traits and require fewer annotations to reach a higher performance. The contrast between target classes and their surroundings contribute greatly to accurate CNN-based mapping (Kattenborn et al., 2020). According to our models’ predictions, distinctive traits could be even more important than the number of annotations for species with constant traits between individuals. Indeed, the abundance of each tree species did not have a major impact on the accuracy of the model despite being strongly unbalanced. The most abundant species, *B. papyrifera*, was not the best classified species, although its classification accuracy was still above average. This species’ features were not particularly distinctive, and they somewhat varied between individuals regarding crown shape and structure and timing of senescence throughout the study site. On the other hand, one of the least abundant species, *L. laricina*, was the second best identified class in the dataset. This species has very distinct features, especially regarding colour and texture. Also, the *L. laricina* crowns were grouped together in a bog, rarely isolated or surrounded by other species. This species often covered most of the surface of the tiles used to train the CNN when it was present. In some cases, underrepresented classes could get crowded by more frequent species in tiles during training (Schiefer et al., 2020). *Larix laricina* was not affected in this way by the unbalanced dataset. A possible reason that less abundant species may have been misclassified could be because they share features with more abundant species (Schiefer et al., 2020). However, the species present in our dataset had few features that overlapped, and the CNN was able to differentiate the less abundant species.

*Pinus strobus* was the class with the highest average F1-score, which is similar to other studies reporting the highest performance for the *Pinus* genus for tree count (Zhang et al., 2022) and crown segmentation and detection (Beloiu et al., 2023; Fricker et al., 2019; Schiefer et al., 2020). A previous study using leaf spectroscopy found that needles of *P. strobus* have a spectral signature that distinguishes this species from other species with high accuracy (Blanchard, 2022). On the imagery, we noticed that *P. strobus* had a distinctive blue-grey colour, similar only to the colour of *L. laricina*, but with a very distinct crown texture. Overall, most of the identification errors for *P. strobus* and other species were between the classes and the background, due to false positives from the model identifying parts of the imagery that have not been annotated or from overestimating the size of the crowns. This indicates that the classes in our dataset have enough distinct features for the CNN to differentiate between each class, especially for classes in different genera.

The major source of misidentification between classes was between species that were closer phylogenetically. Species in the same genus may share features and have a similar appearance in the imagery (Onishi et al., 2022), making them more prone to misclassifications by the CNN. This was especially true for the three maple species, *A. pensylvanicum*, *A. rubrum* and *A. saccharum*, which were often misclassified among each other throughout the growing season. Also, trees of the same functional groups and growing in similar environments may have a similar texture and light acquisition performance (Onishi et al., 2022). Due to *A. rubrum*’s plasticity (Royer et al., 2008, 2009), its functional traits such as leaf shape vary depending on the conditions in which it grows, making it resemble *A. saccharum* and sometimes *A. pensylvanicum*. Along with its variability during senescence, this variability in *A. rubrum*’s appearance would explain the lower F1-score despite the high abundance of annotations for this class. While the variability of *A. rubrum*’s features increased the proportion of misclassification, its high abundance within the dataset seems to mitigate these effects since it was overall well identified.

Our study shows the effect of phenology, specifically autumn leaf senescence, on the performance of a CNN in identifying and segmenting tree crowns in a temperate forest. Contrary to our hypothesis, in October, when almost all leaves of tree species at the top of the canopy had changed colours, the CNN produced the least accurate predictions. We attribute this to the important leaf fall and the intra and interspecific variability of the timing of the different events within senescence. Nevertheless, our results show that the predictions produced by a CNN remained relatively accurate all throughout the growing season, from the end of spring to autumn. While this offers an insight into the way leaf senescence may influence a CNN, imagery with a higher temporal resolution would provide more detailed information on the senescence process in this temperate forest, especially for species with highly asynchronous timing. This can be important when it comes to monitoring temperate forest using a UAV and deep learning, for phenology monitoring or tree mapping.

## 5. Conclusion

Understanding the influence of leaf phenology on the capacity of a CNN to accurately identify and segment tree crowns is crucial to improving and optimizing our use of UAVs and remote sensing in forests. High spatial and temporal resolution imagery can help to understand and map these ecosystems, and by leveraging the phenological traits associated with temperate forests, it may help to understand these complex processes. In this study, we show how autumn phenology, specifically leaf senescence in a temperate mixed forest, can affect the performance of a CNN in segmenting individual tree crowns and identifying them to the species level. While peak colouring can be beneficial for some deciduous species, for most species, it is detrimental due to asynchronous leaf fall, especially in a heterogenous forest. Earlier in the season, at the start of leaf colouring, our models gave the best predictions overall. This further highlights the importance of the timing of imagery acquisition for optimized conditions for species mapping, a factor mentioned by others (Hill et al., 2010). The models presented in this study could be optimized by acquiring imagery at a higher temporal resolution, especially during leaf senescence, and to include earlier spring phenology to create a more complete representation of the growing season. To account for the variability of the field site, integrating topo-edaphic variables could allow for a better understanding of the classification. More recent deep learning models could also give better predictions, but the objective of the study was to compare the performance of the models to one another, not to achieve the best predictions.

## Acknowledgments

The authors thank Noémie Lacombe, Ariane Roberge, Antoine Caron-Guay and Étienne Morissette whose contribution to the field work and the annotations were essential to building the dataset used in this study. The authors also thank the staff at the Station de biologie des Laurentides for the lodgings during the field work and access to the field site and equipment. This study was funded by the Canada First Research Excellence Fund to the Institute for Data Valorisation (IVADO, Grant number PRF-2021-02), a Discovery grant from the Natural Science and Engineering Research Council of Canada to E.L. (NSERC, Grant number RGPIN-2019-04537) and from the Canada Research Chair in Plant Functional Biodiversity to E.L. (Grant number 950-231858).

