## Supplementary Material for "Influence of Temperate Forest Autumn Leaf Phenology on Segmentation of Tree Species from UAV Imagery Using Deep Learning"

### Supplementary Materials

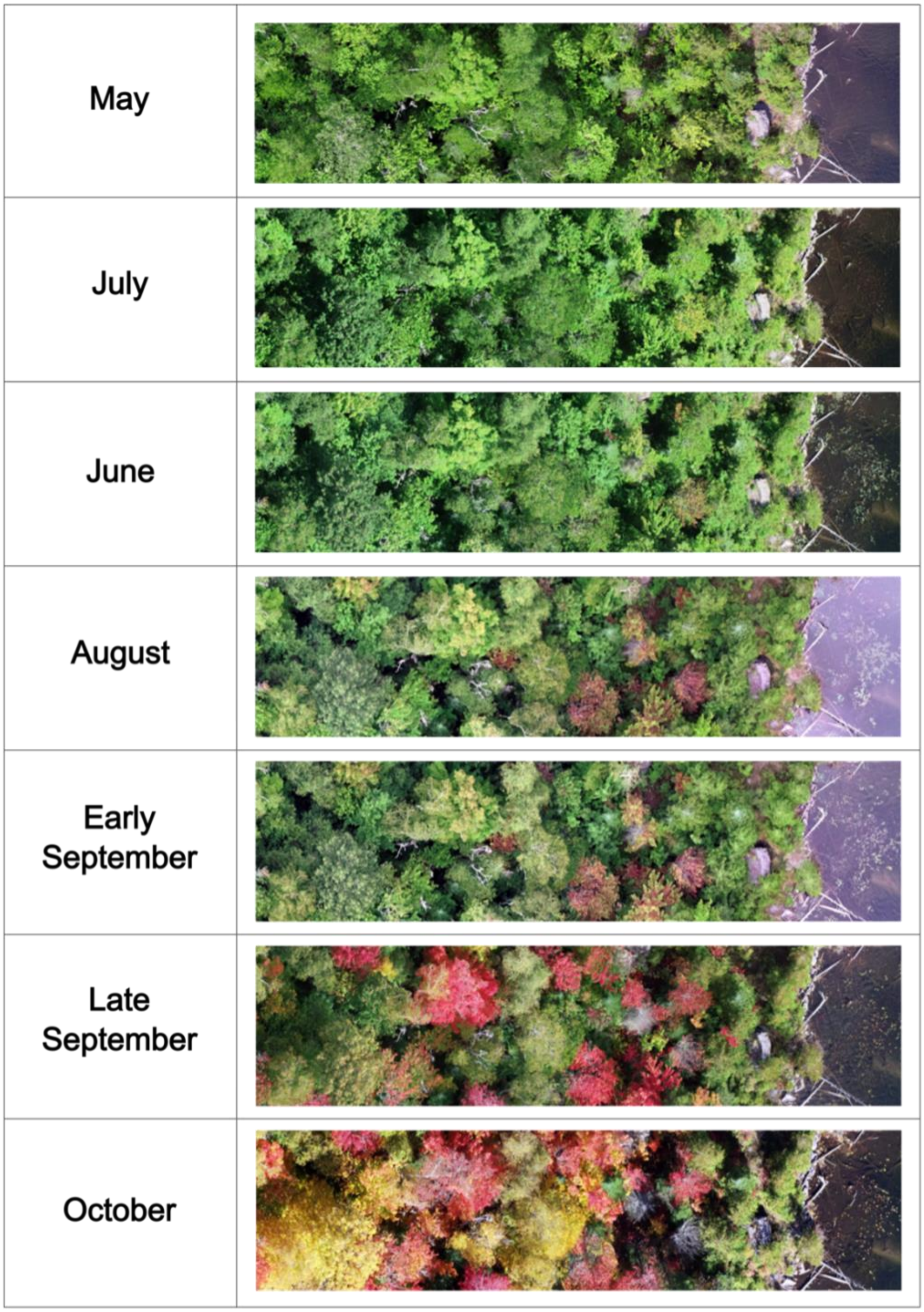


**Figure S1 Illumination conditions for each date of acquisition**
The imagery with more diffuse light conditions are from August and early September, and the imagery with more sunny light conditions are from June, July, and October. May and late September have intermediate light conditions.

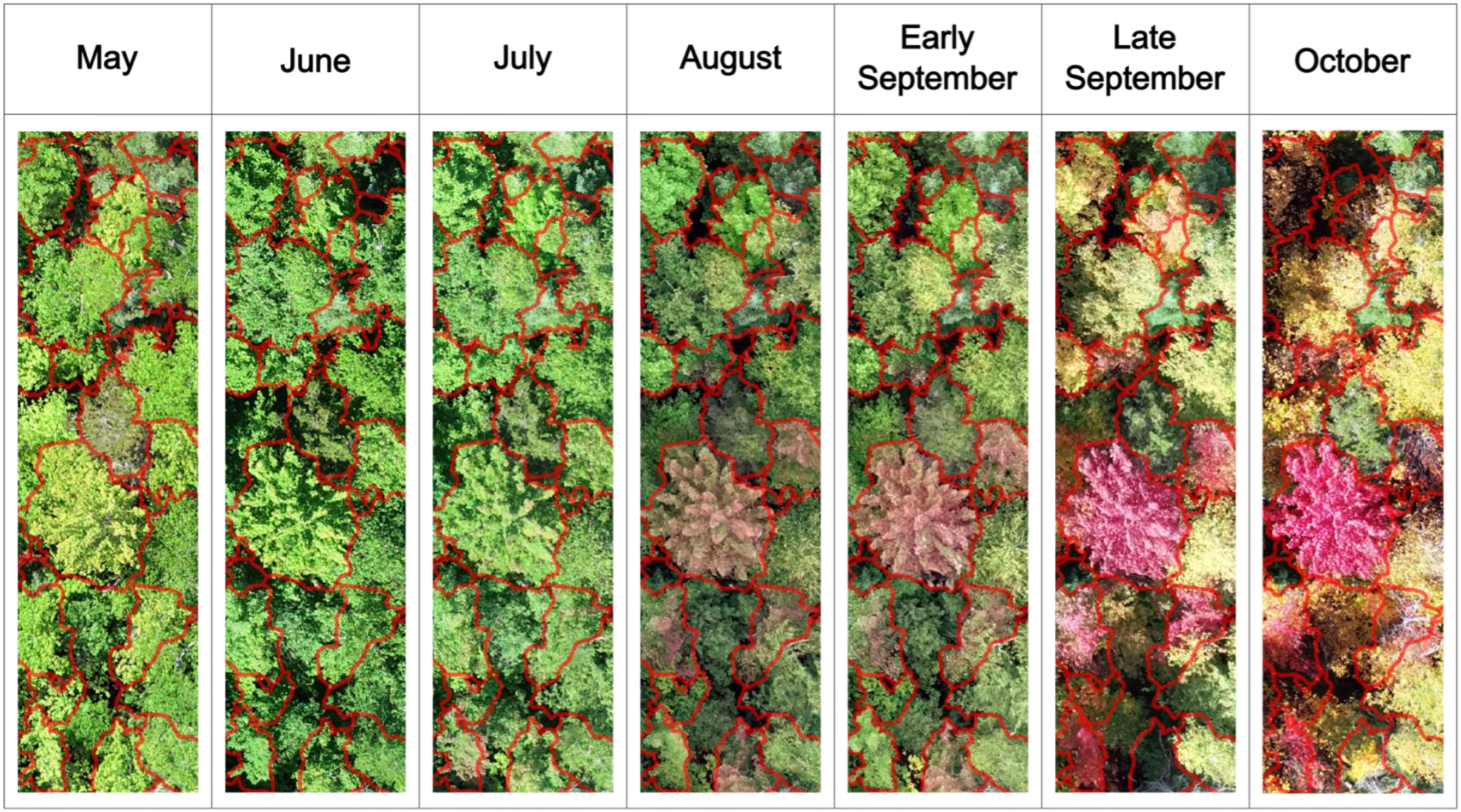


**Figure S2 Example of annotated tree crowns**
It is possible to see that the annotations (contour of the polygons in red) are well aligned with the orthomosaic for each date, making it possible to apply the segmentation done on the imagery of early September onto the orthomosaics from other dates.

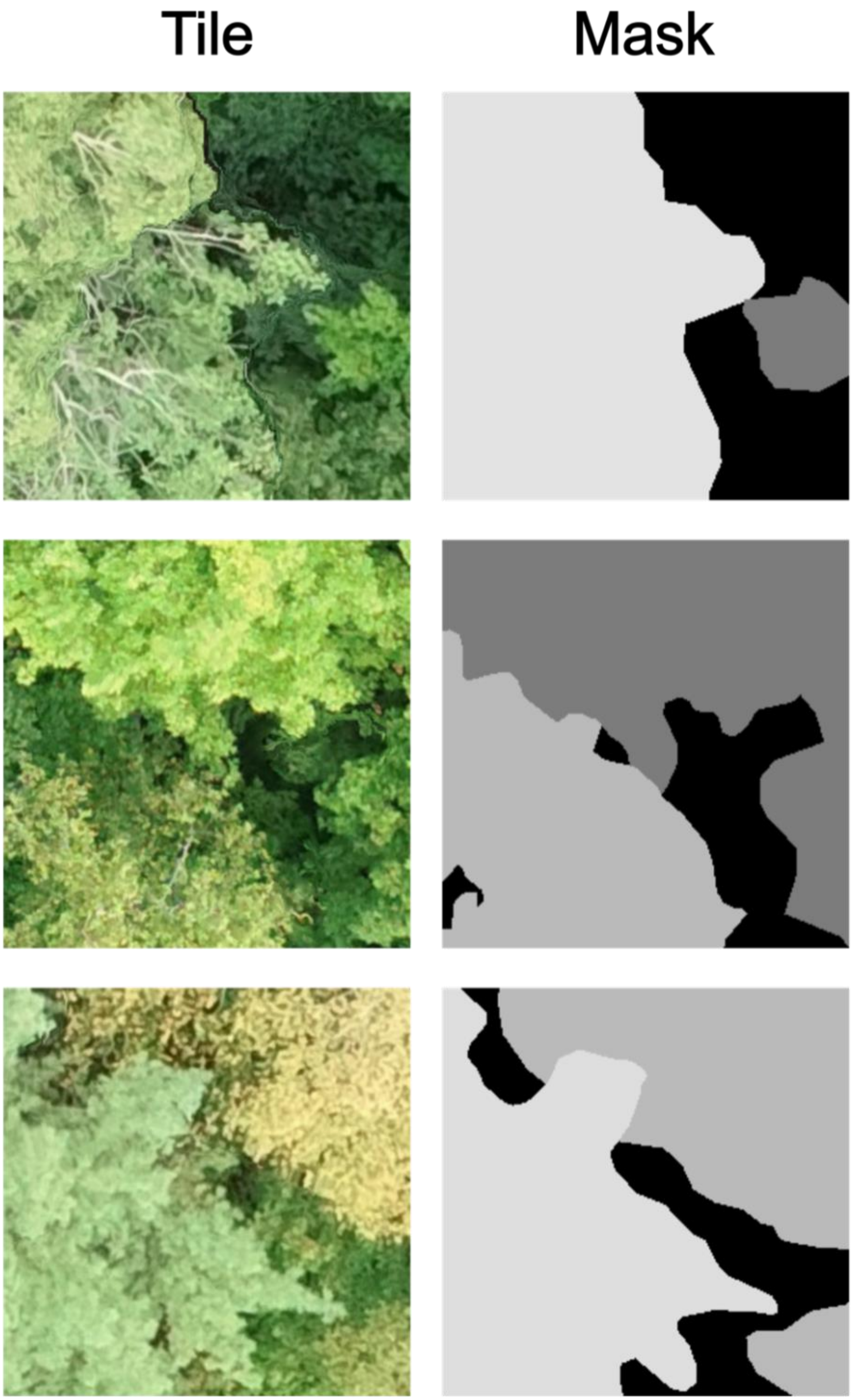


**Figure S3 Example of tiles and masks from the early September imagery**
The tiles represent a section of the orthomosaic of 256 x 256 pixels and the masks represent the annotations for that section in raster format. The different colours in the masks correspond to different classes.

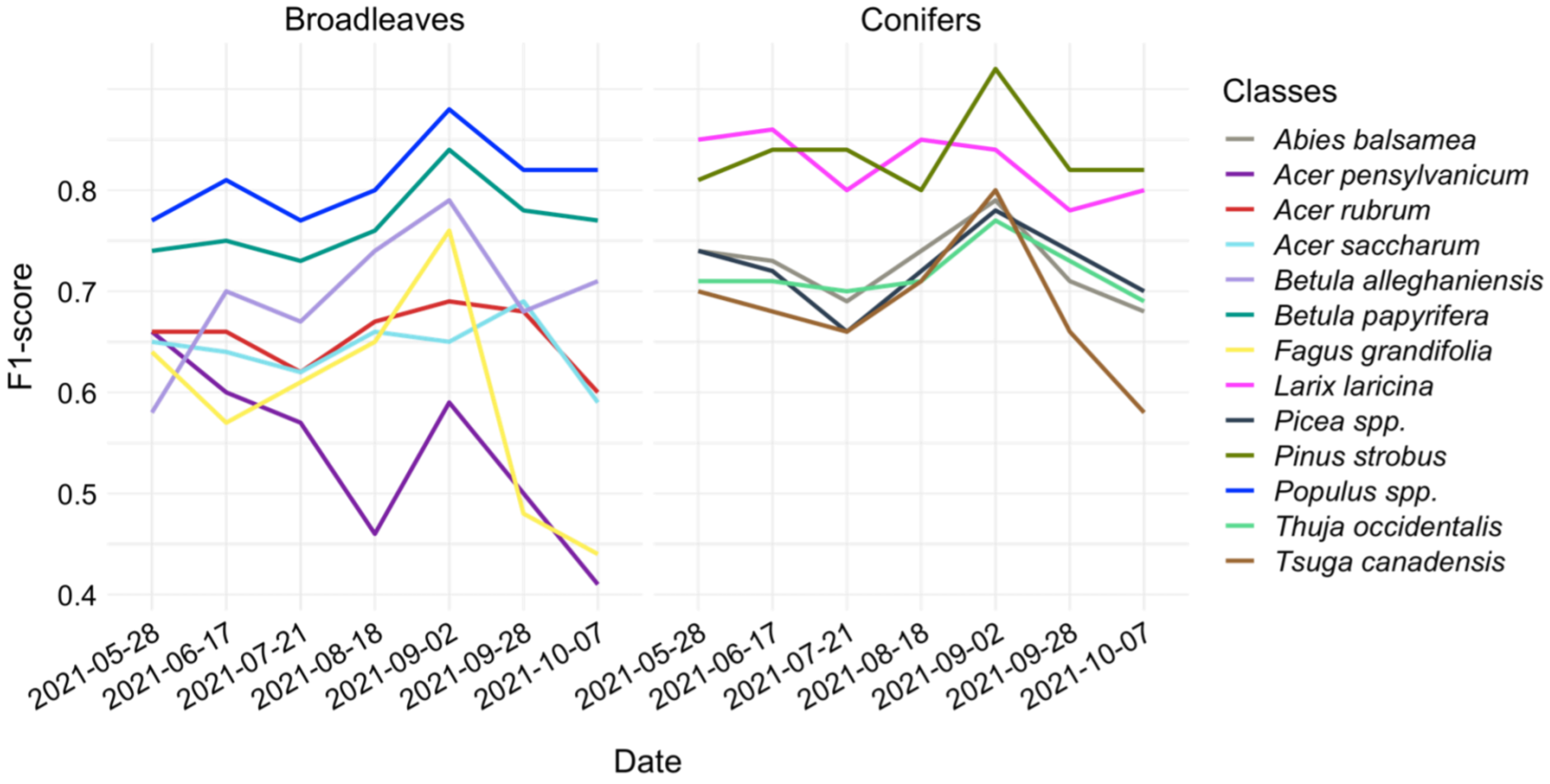


**Figure S4 Variation of the F1-scores for each class for the seven dates**
The temporal variation of the F1-score throughout the growing season is more variable for deciduous broadleaf species (left) than it is for conifers (right). Most species show a peak in their F1-score for the September 2$^{nd}$ model, and often a decrease in the F1-score for the October 7$^{th}$ model.

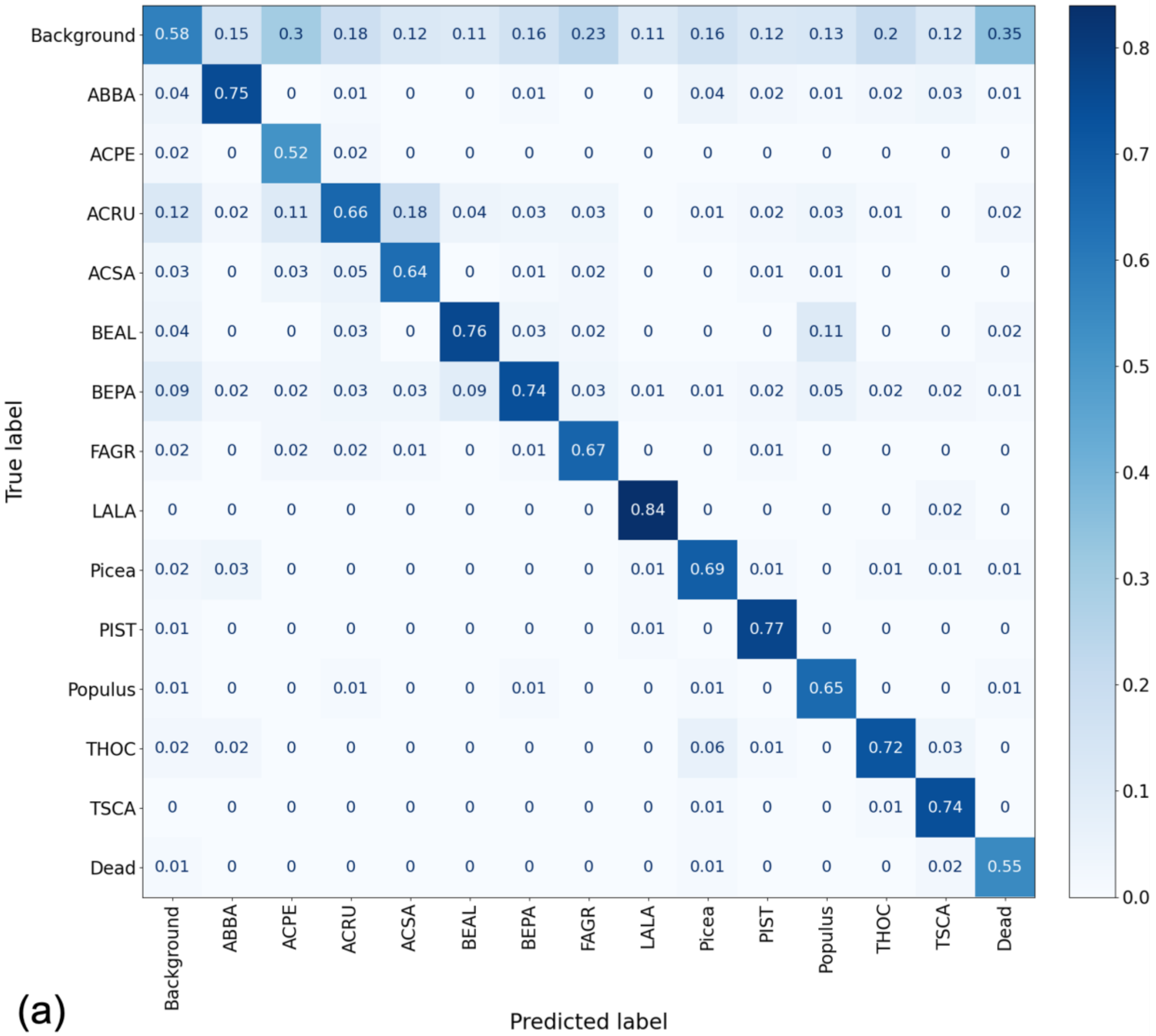
True label
Predicted label
Background ABBA ACPE ACRU ACSA BEAL BEPA FAGR LALA Picea PIST Populus THOC TSCA Dead
Background 0.58 0.15 0.3 0.18 0.12 0.11 0.16 0.23 0.11 0.16 0.12 0.13 0.2 0.12 0.35
ABBA 0.04 0.75 0 0.01 0 0 0.01 0 0 0.04 0.02 0.01 0.02 0.03 0.01
ACPE 0.02 0 0.52 0.02 0 0 0 0 0 0 0 0 0 0 0
ACRU 0.12 0.02 0.11 0.66 0.18 0.04 0.03 0.03 0 0.01 0.02 0.03 0.01 0 0.02
ACSA 0.03 0 0.03 0.05 0.64 0 0.01 0.02 0 0 0.01 0.01 0 0 0
BEAL 0.04 0 0 0.03 0 0.76 0.03 0.02 0 0 0 0.11 0 0 0.02
BEPA 0.09 0.02 0.02 0.03 0.03 0.09 0.74 0.03 0.01 0.01 0.02 0.05 0.02 0.02 0.01
FAGR 0.02 0 0.02 0.02 0.01 0 0.01 0.67 0 0 0.01 0 0 0 0
LALA 0 0 0 0 0 0 0 0 0.84 0 0 0 0 0.02 0
Picea 0.02 0.03 0 0 0 0 0 0 0.01 0.69 0.01 0 0.01 0.01 0.01
PIST 0.01 0 0 0 0 0 0 0 0.01 0 0.77 0 0 0 0
Populus 0.01 0 0 0.01 0 0 0.01 0 0 0.01 0 0.65 0 0 0.01
THOC 0.02 0.02 0 0 0 0 0 0 0 0.06 0.01 0 0.72 0.03 0
TSCA 0 0 0 0 0 0 0 0 0 0.01 0 0 0.01 0.74 0
Dead 0.01 0 0 0 0 0 0 0 0 0.01 0 0 0 0.02 0.55
0.0
0.1
0.2
0.3
0.4
0.5
0.6
0.7
0.8

(a)

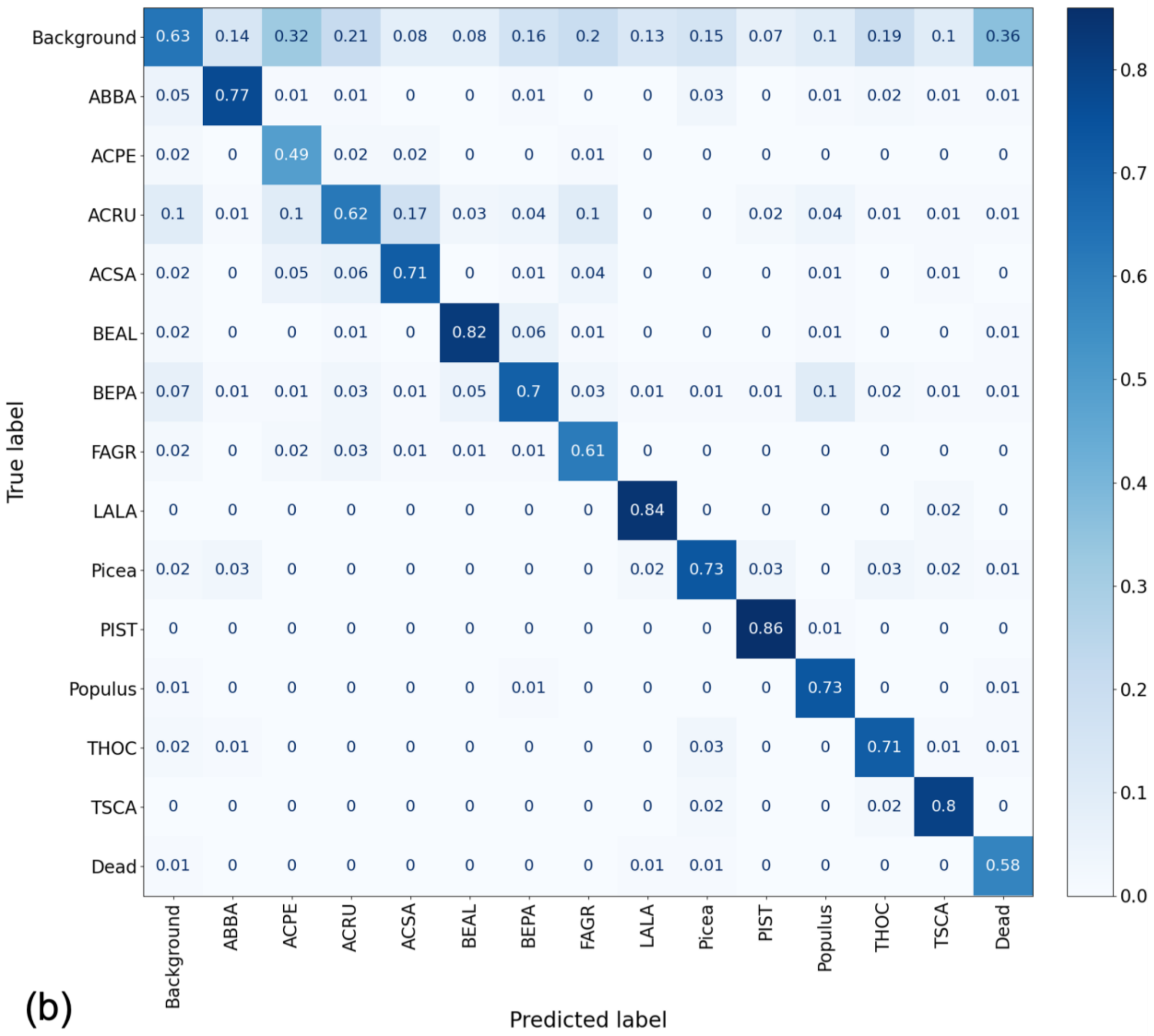
True label
Predicted label
Background ABBA ACPE ACRU ACSA BEAL BEPA FAGR LALA Picea PIST Populus THOC TSCA Dead
Background 0.63 0.14 0.32 0.21 0.08 0.08 0.16 0.2 0.13 0.15 0.07 0.1 0.19 0.1 0.36
ABBA 0.05 0.77 0.01 0.01 0 0 0.01 0 0 0.03 0 0.01 0.02 0.01 0.01
ACPE 0.02 0 0.49 0.02 0.02 0 0 0.01 0 0 0 0 0 0 0
ACRU 0.1 0.01 0.1 0.62 0.17 0.03 0.04 0.1 0 0 0.02 0.04 0.01 0.01 0.01
ACSA 0.02 0 0.05 0.06 0.71 0 0.01 0.04 0 0 0 0.01 0 0.01 0
BEAL 0.02 0 0 0.01 0 0.82 0.06 0.01 0 0 0 0.01 0 0 0.01
BEPA 0.07 0.01 0.01 0.03 0.01 0.05 0.7 0.03 0.01 0.01 0.01 0.1 0.02 0.01 0.01
FAGR 0.02 0 0.02 0.03 0.01 0.01 0.01 0.61 0 0 0 0 0 0 0
LALA 0 0 0 0 0 0 0 0 0.84 0 0 0 0 0.02 0
Picea 0.02 0.03 0 0 0 0 0 0 0.02 0.73 0.03 0 0.03 0.02 0.01
PIST 0 0 0 0 0 0 0 0 0 0 0.86 0.01 0 0 0
Populus 0.01 0 0 0 0 0 0.01 0 0 0 0 0.73 0 0 0.01
THOC 0.02 0.01 0 0 0 0 0 0 0 0.03 0 0 0.71 0.01 0.01
TSCA 0 0 0 0 0 0 0 0 0 0.02 0 0 0.02 0.8 0
Dead 0.01 0 0 0 0 0 0 0 0.01 0.01 0 0 0 0 0.58
0.0 0.1 0.2 0.3 0.4 0.5 0.6 0.7 0.8
(b)

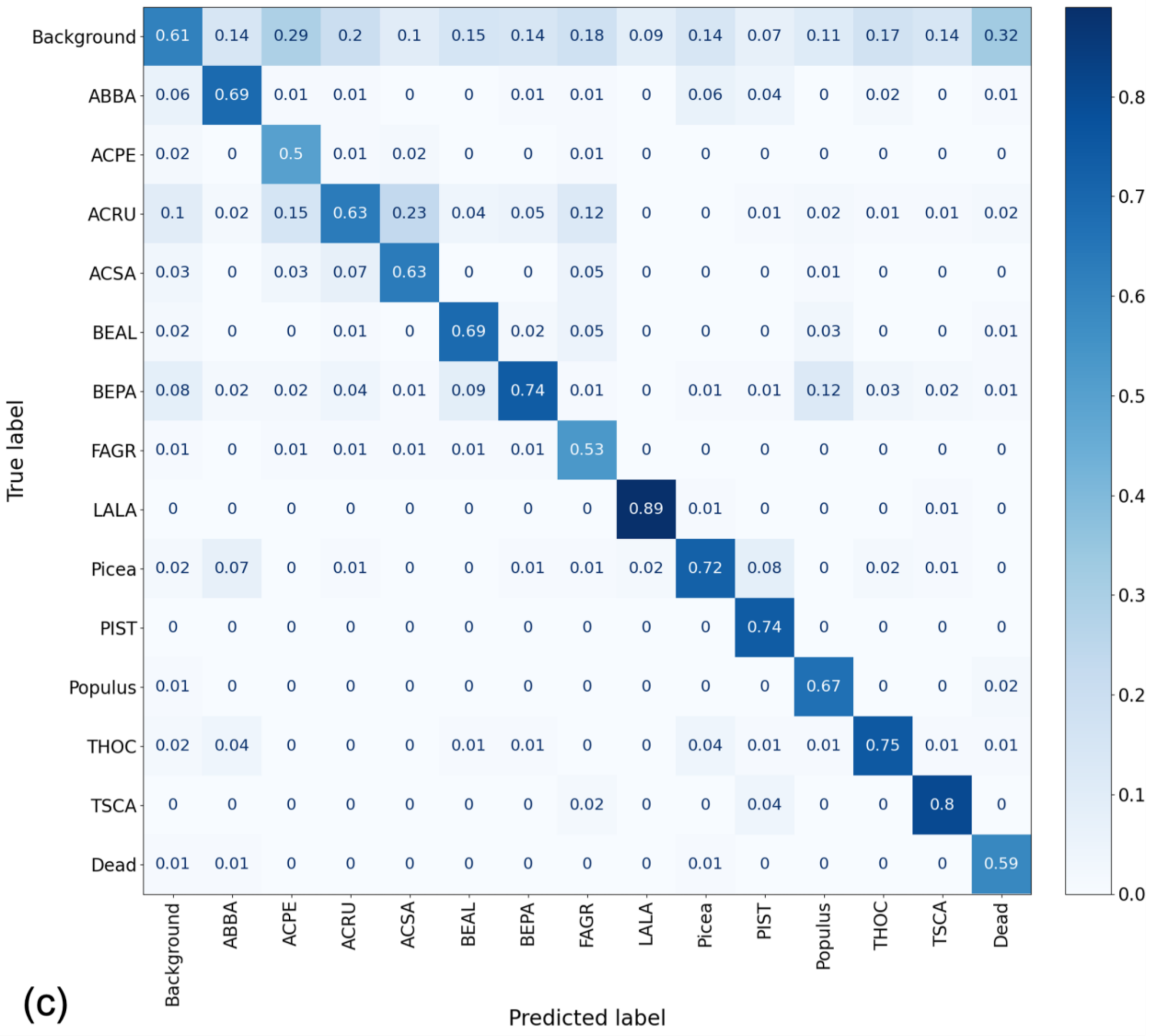

True label
Predicted label
Background 0.61 0.14 0.29 0.2 0.1 0.15 0.14 0.18 0.09 0.14 0.07 0.11 0.17 0.14 0.32
ABBA 0.06 0.69 0.01 0.01 0 0 0.01 0.01 0 0.06 0.04 0 0.02 0 0.01
ACPE 0.02 0 0.5 0.01 0.02 0 0 0.01 0 0 0 0 0 0 0
ACRU 0.1 0.02 0.15 0.63 0.23 0.04 0.05 0.12 0 0 0.01 0.02 0.01 0.01 0.02
ACSA 0.03 0 0.03 0.07 0.63 0 0 0.05 0 0 0 0.01 0 0 0
BEAL 0.02 0 0 0.01 0 0.69 0.02 0.05 0 0 0 0.03 0 0 0.01
BEPA 0.08 0.02 0.02 0.04 0.01 0.09 0.74 0.01 0 0.01 0.01 0.12 0.03 0.02 0.01
FAGR 0.01 0 0.01 0.01 0.01 0.01 0.01 0.53 0 0 0 0 0 0 0
LALA 0 0 0 0 0 0 0 0 0.89 0.01 0 0 0 0.01 0
Picea 0.02 0.07 0 0.01 0 0 0.01 0.01 0.02 0.72 0.08 0 0.02 0.01 0
PIST 0 0 0 0 0 0 0 0 0 0 0.74 0 0 0 0
Populus 0.01 0 0 0 0 0 0 0 0 0 0 0.67 0 0 0.02
THOC 0.02 0.04 0 0 0 0.01 0.01 0 0 0.04 0.01 0.01 0.75 0.01 0.01
TSCA 0 0 0 0 0 0 0 0.02 0 0 0.04 0 0 0.8 0
Dead 0.01 0.01 0 0 0 0 0 0 0 0.01 0 0 0 0 0.59
Background ABBA ACPE ACRU ACSA BEAL BEPA FAGR LALA Picea PIST Populus THOC TSCA Dead
0.0 0.1 0.2 0.3 0.4 0.5 0.6 0.7 0.8
(c)

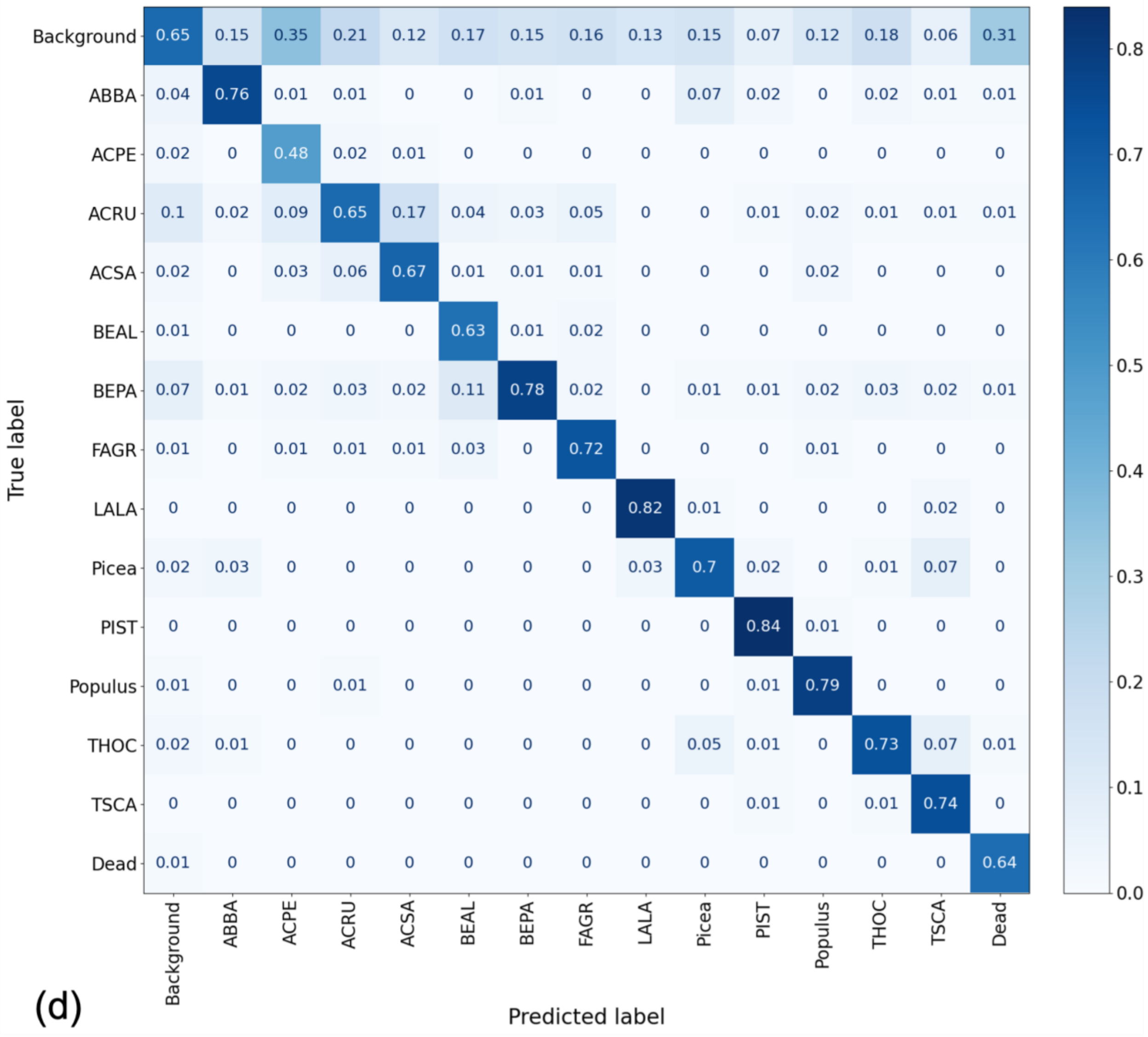

True label
Predicted label
Background
ABBA
ACPE
ACRU
ACSA
BEAL
BEPA
FAGR
LALA
Picea
PIST
Populus
THOC
TSCA
Dead
(d)

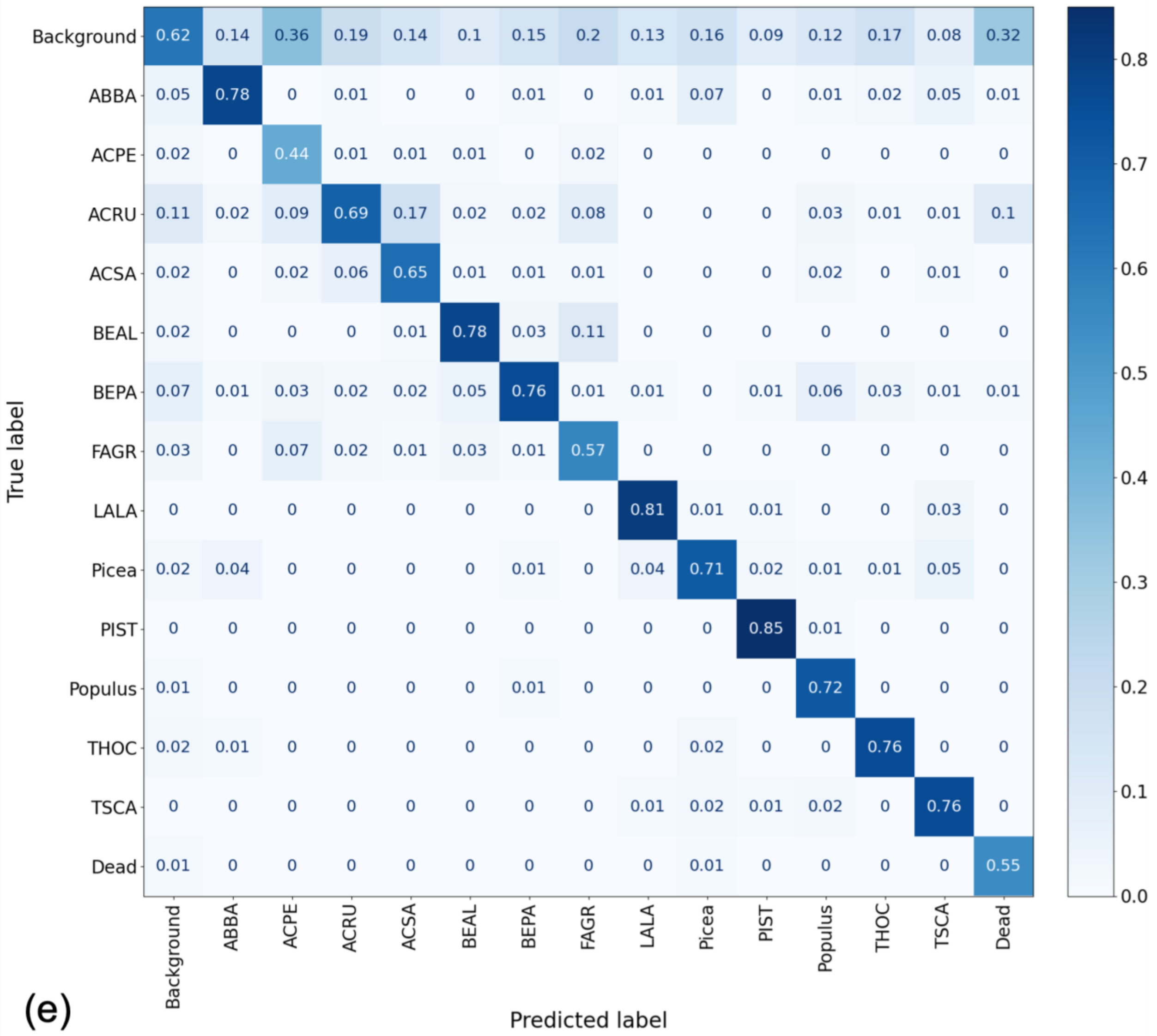


**Figure S5** **Confusion matrices for each class for the remaining five dates**

The values represent the overlap (from 0-1) between the annotations (true label) and the predictions (predicted label) done by the model on unseen imagery from (a) May 28th, 2021, (b) June 17th, 2021, (c) July 21st, 2021, (d) August 18th, 2021, (e) September 28th, 2021. ABBA: *Abies balsamea*, ACPE: *Acer pensylvanicum*, ACRU: *Acer rubrum*, ACSA: *Acer saccharum*, BEAL: *Betula alleghaniensis*, BEPA: *Betula papyrifera*, FAGR: *Fagus grandifolia*, LALA: *Larix laricina*, Picea: *Picea spp.*, PIST: *Pinus strobus*, Populus: *Populus spp.*, THOC: *Thuja occidentalis*, TSCA: *Tsuga canadensis*.
